# Event boundaries drive norepinephrine release and distinctive neural representations of space in the rodent hippocampus

**DOI:** 10.1101/2024.07.30.605900

**Authors:** Sam McKenzie, Alexandra L. Sommer, Tia N. Donaldson, Infania Pimentel, Meenakshi Kakani, Irene Jungyeon Choi, Ehren L. Newman, Daniel F. English

**Affiliations:** Department of Neurosciences, University of New Mexico Health Science Center, Albuquerque, NM 87106; Department of Mechanical Engineering, Tufts School of Engineering, Medford MA 02155; Department of Biology, Virginia Commonwealth University, Richmond, VA 23284; Psychological and Brain Sciences, Indiana University, Bloomington, IN, 47405; Program in Neuroscience, Indiana University, Bloomington, IN, 47405; School of Neuroscience, Virginia Tech, Blacksburg, VA, 24061

## Abstract

Episodic memories are temporally segmented around event boundaries that tend to coincide with moments of environmental change. During these times, the state of the brain should change rapidly, or reset, to ensure that the information encountered before and after an event boundary is encoded in different neuronal populations. Norepinephrine (NE) is thought to facilitate this network reorganization. However, it is unknown whether event boundaries drive NE release in the hippocampus and, if so, how NE release relates to changes in hippocampal firing patterns. The advent of the new GRAB_NE_ sensor now allows for the measurement of NE binding with sub-second resolution. Using this tool in mice, we tested whether NE is released into the dorsal hippocampus during event boundaries defined by unexpected transitions between spatial contexts and presentations of novel objections. We found that NE binding dynamics were well explained by the time elapsed after each of these environmental changes, and were not related to conditioned behaviors, exploratory bouts of movement, or reward. Familiarity with a spatial context accelerated the rate in which phasic NE binding decayed to baseline. Knowing when NE is elevated, we tested how hippocampal coding of space differs during these moments. Immediately after context transitions we observed relatively unique patterns of neural spiking which settled into a modal state at a similar rate in which NE returned to baseline. These results are consistent with a model wherein NE release drives hippocampal representations away from a steady-state attractor. We hypothesize that the distinctive neural codes observed after each event boundary may facilitate long-term memory and contribute to the neural basis for the primacy effect.

## Introduction

Determining neurobiological mechanisms by which the hippocampus supports the formation of memories for distinct episodes remains a major outstanding challenge. Norepinephrine (NE) signaling is hypothesized to play a key role in organizing memory into episodes demarcated by event boundaries^1^. Yet, the situations in which NE is released in the hippocampus, and the effects of NE on hippocampal coding, are not well understood. Here, we use the GRAB_NE_ sensor and analysis of neuronal spiking dynamics to test the hypothesis that NE release occurs at event boundaries and aligns with changes in neural coding that promote long-term memory.

Prior work suggests that NE release from the locus coeruleus (LC) may facilitate event segmentation by modulating the induction threshold for synaptic plasticity^2–11^, facilitating reorganization of which neurons are active before and after unexpected salient events^12^, and changing how neurons encode their environment at the time of transmitter release^13^. NE release from the LC causes immediate changes in the excitability and activity of neurons across the hippocampal formation^14–23^. Electrical stimulation of the LC acutely silences most hippocampal neurons^24,25^ while simultaneously increasing firing in the subset of neurons that respond to salient stimuli^25^, an observation that motivated the hypothesis that NE sets the gain of the neuronal input/output curve^26^. Computational models predict that NE-induced changes in gain should promote network shifts by lowering the activation energy for transitioning between learned states/attractors^27–32^. Hippocampal place fields remap^33^ (change place field position), with learning^34–38^ and also after salient changes in an animal’s environment^39–41^, offering an attractive correlate to assess LC-induced reset^42^.

NE also facilitates synaptic plasticity^2^. Plasticity-related signaling is needed for the reactivation of waking spiking activity during subsequent sharp-wave ripples replay events^34,43,44^. Neuronal replay is important for memory consolidation^45^ and variations in the content of replay may dictate which moments are remembered and which are forgotten^46^. Stimulation of dopaminergic terminals from the ventral midbrain^47^, as well as natural reward^48^, enhances synaptic plasticity and can promote reactivation. It is unknown whether moments of elevated noradrenergic release similarly bias subsequent replay, though such a relationship has been predicted^49^.

Micro-dialysis studies have revealed that NE is released in the hippocampus after exposure to novel environments^50,51^, physical restraint/handling^50,51^, or after exposure to novel combinations of familiar objects^52^. This method samples average NE concentration over a minutes-long period and therefore cannot resolve whether release is related to the experimental stimuli or the behaviors associated with those stimuli; for example, mice move more in novel spaces. The low sampling resolution of micro-dialysis also precludes relating moment-to-moment changes in neural coding with fluctuations in NE concentration. Using the recently developed GRAB_NE_ sensor^53^, which can measure NE release with sub-second resolution, hippocampal NE levels were shown to increase immediately after delivering an electrical shock and decrease around freezing^54^. This pattern could indicate a relationship between NE around encoding and retrieval events, or alternatively, may arise due to a relationship between NE release and overall levels of movement or arousal, which in this case co-varied with different phases of the experiment. In support of this latter interpretation, a previous study found that the firing rate of LC neurons positively correlates with acceleration^55^. Others have reported that LC neurons fire in response to unexpected salient stimuli^56–63^, including reward prediction errors^64–66^. Such surprise-related activity of LC neurons may cause NE release at the moments when event boundaries are thought to occur, however, such a relationship is not guaranteed as NE release is also modulated at the level of the axon terminal^67^.

To better understand how hippocampal NE release dynamics relate to event boundaries and the associated neuronal response, we used the GRAB_NE_ sensor to examine how NE release is related to event boundaries imposed by unexpected transitions between testing environments and the introduction of novel objects. We also tested how these signals are affected by moment-to-moment fluctuations in behavior and reward availability, and how NE release dynamics change over the course of learning. Knowing when NE is expressed, we then assessed whether these moments are associated with changes in neural coding as predicted by prominent models of NE function. Our findings support a model in which NE release around event boundaries scales with the deviance between current and previously stored neural representations.

## Results

To investigate the dynamics of NE release and binding in the dorsal hippocampus, the GRAB_NE_ genetically encoded fluorescent indicator^53^ was virally delivered to dorsal CA1 (Figure 1A), and optic fibers were chronically implanted in C57BL6/J mice (N = 8 mice, N = 3 female) unilaterally targeting the injection site. The main dependent measure was the emission intensity of the NE-derived signal (experimental excitation λ = 465-nm) with corrections for mechanical instability (isosbestic excitation λ = 405-nm) and photobleaching, and normalized by the mean and standard deviation recorded during a 10-minute homecage baseline (see Methods); this measurement will be referred to as Signal_NE_. The Signal_NE_ derived from the GRAB_NE_ sensor was validated in our hands by showing that the noradrenergic reuptake inhibitor desipramine caused a significant increase in Signal_NE_ relative to vehicle injections (Figure S1A). Likewise, noradrenergic α2 receptor antagonism with yohimbine (from which GRAB_NE_ was derived) disrupted normally strong Signal_NE_ (Figure S1B).

**Figure 1.**
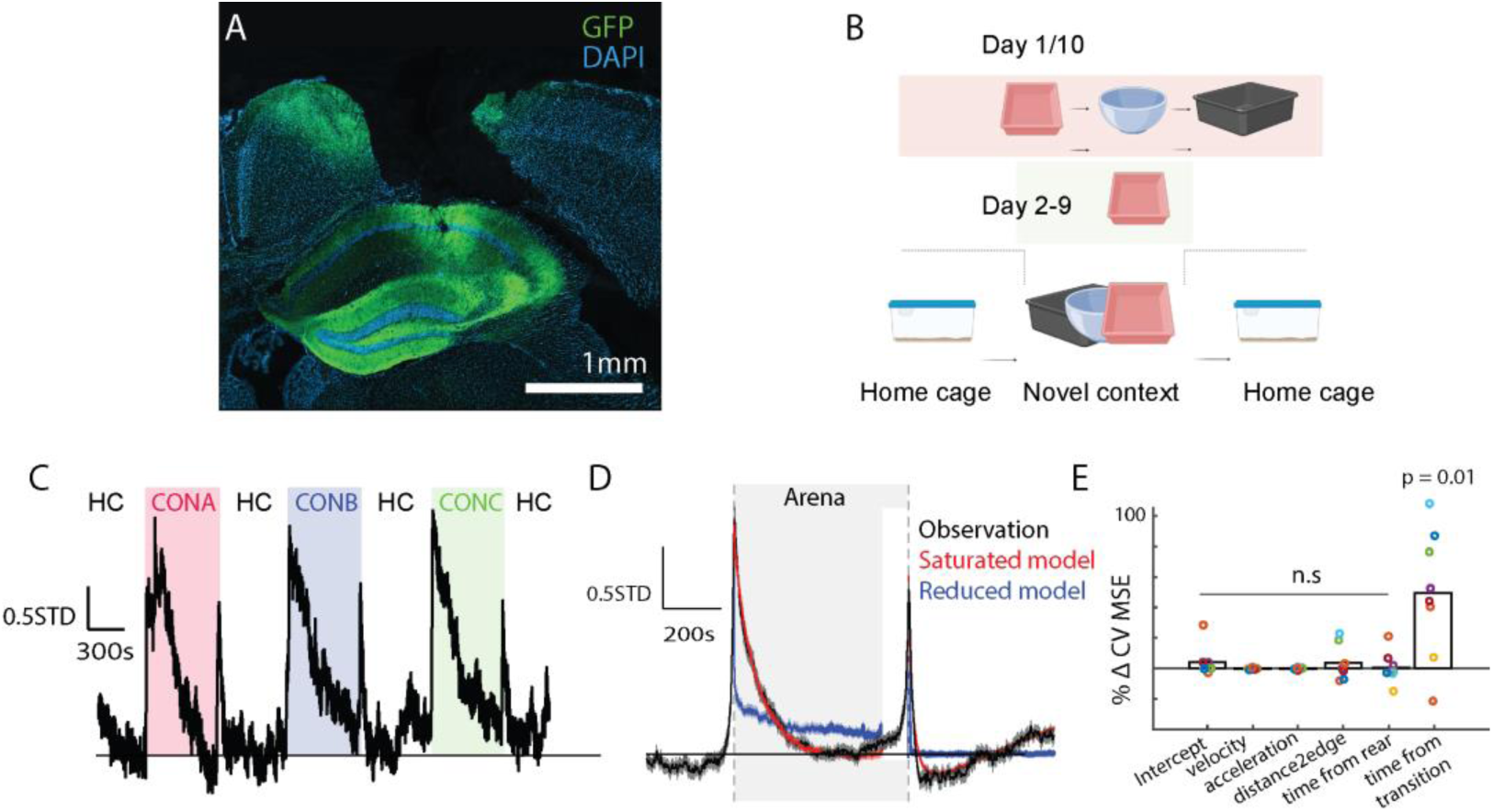
Time from context transition controls Signal_NE_ when mice are moved to novel arenas. A) Histological confirmation of GRAB_NE_ expression (GFP) and fiber placement over dorsal CA1. B) Schematic of experimental timeline. C) Example session showing increases in Signal_NE_ around each context and homecage (HC) transition. D) Mean Signal_NE_ measured across all transitions (black) and cross-validated prediction from the saturated model (red) or a reduced model lacking terms related to time from transfer (blue). E) Change in CVMSE due to removal of various potential behavioral variables. Only removal of the terms related to time from transition significantly decreased model performance (*t(7) = 3.30, p = 0.01*).

### Signal_NE_ exponentially decays after transfer to a novel arena

Moving between environments causes a large reorganization in which hippocampal neurons are active^33,68^. To test how NE release relates to this cause of network reorganization, Signal_NE_ was measured as mice were transferred from their home cage to a novel testing arena that, over days, became more familiar to the subject (Figure 1B,C). Averaging across all exposures, Signal_NE_ increased immediately upon entry to the testing arena and exponentially decayed to a steady state over minutes (Figure 1D). The NE dynamics may be related to the transition itself, or the incidence of behaviors that occur immediately following exposure to an unfamiliar space. For example, in the moments after transition, mice tended to spend more time close to the edges of the environment (thigmotaxis) and tended to move more rapidly (Figure S1C). We quantified how NE release relates to five potential behavioral covariates: time from arena entry, acceleration, velocity, distance from edge, and time from rearing. These five behavioral variables were themselves correlated (Figure S1D). Univariate analysis revealed strong, positive co-modulation of Signal_NE_ with acceleration (*t(7) = 4.54, p = 0.002*) and modest positive correlation with velocity (*t(7) = 2.32, p = 0.05*)(Figure S1E). Signal_NE_ also correlated with distance to the edge of the environment (*t(7) = -2.37, p = 0.05*)(Figure S1E,F), and showed transient changes around rearing events in a subset of animals (Figure S1E,G). Such covariation in putative factors driving NE release complicates assessment of whether NE release dynamics relate to the contextual transition *per se*, or whether NE is more closely associated with novelty-related behaviors. The sub-second temporal resolution of the GRAB_NE_ sensors allows disambiguation of these scenarios.

To identify the independent variable with the greatest explanatory power, we performed backward stepwise regression on a non-linear model defined by the five behavioral variables of interest. Time from transition was modeled with two terms: a positive term with a fast decay and a negative term with a slower decay to capture decreases in NE observed after some transitions. Cross-validated mean squared error (CVMSE) was calculated for the full, saturated model and for a reduced model in which one of the five variables (or the intercept) has been dropped. Significant decreases in model fit were only observed after dropping the time from context entry independent variable (Figure 1E). Despite apparent modulation of Signal_NE_ with movement, the critical factor in predicting Signal_NE_ was the time from event transition.

### Signal_NE_ exponentially decays after transfer to a familiar linear track

LC activity and NE release have been related to reward^65,69^ and acceleration^55^. The physical dimensions of the testing arenas prohibited moments of high acceleration or velocity and the recording sessions lacked appetitive reward conditions. We therefore sought to test whether Signal_NE_ was under the control of event boundary transitions even when mice engaged in a learned task in which subjects must run to receive water reward on a linear track, a standard apparatus for studying hippocampal physiology.

Here, we considered five independent variables: time from linear track entry, acceleration, velocity, distance from the edge of the track, and time from reward. As was observed in the novel arena experiments, NE increased rapidly upon entry to the linear track and decayed to a steady state (Figure 2 A,B). Hippocampal NE was not modulated around reward delivery (signaled with an audible solenoid click) nor movement (Figure S2). The stepwise regression analysis showed that removing time from entry, but no other term, significantly decreased our ability to predict fluctuations in Signal_NE_ (Figure 2C). These results show that, even in the context of an appetitive task that requires conditioned responses, time from transition is the dominant factor in explaining hippocampal NE release.

**Figure 2.**
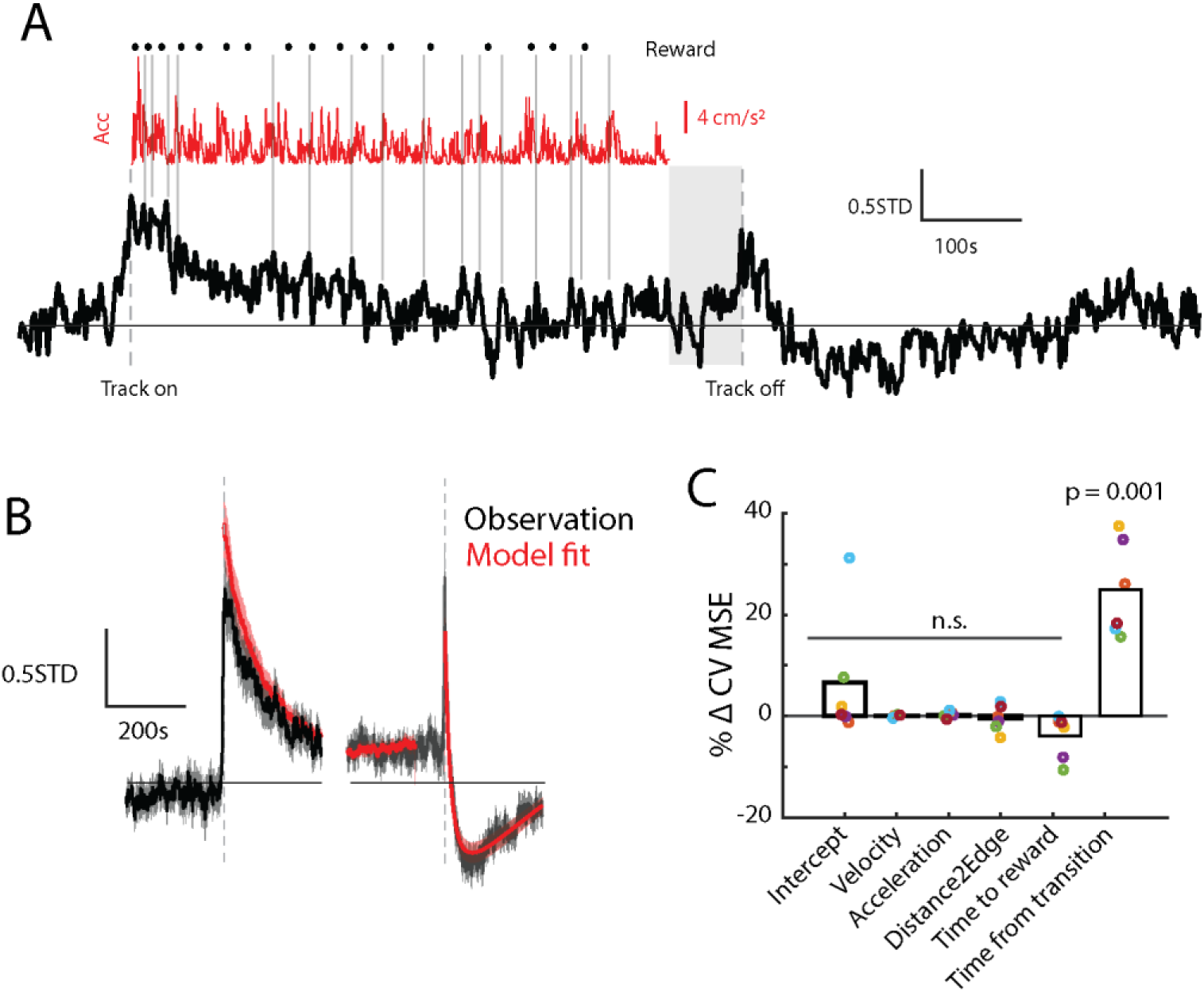
Time from context transition controls Signal_NE_ when mice are moved to a linear track. A) Example session showing Signal_NE_ (black) aligned with acceleration (red) and reward delivery (**·**). Vertical gray lines show that local peaks in Signal_NE_ do not align to bouts of acceleration nor reward timing. Shaded area shows last 60s before removing from track during which Signal_NE_ was not modeled. B) Mean Signal_NE_ measured across all linear track transitions (black) and cross-validated prediction from the saturated model (red). C) Change in CVMSE due to removal of various potential behavioral variables. Only removal of the terms related to time from transition significantly decreased model performance (*t(7) = 7.20,p = 0.0008*).

### Signal_NE_ exponentially decays after introduction of a novel object

In experiments that have studied event boundaries in people, the modality of the information is often non-spatial (e.g. the color of a picture background^1,70^) and LC firing has been related to object sampling in the rodent^62^. Therefore, we tested whether the introduction of a novel object could likewise signal an event boundary to the mouse that would be associated with a transient increase in Signal_NE_.

Five novel objects were consecutively introduced to the mouse, each for five minutes starting 10 minutes after the mouse was transferred to a familiar arena, a timeline designed to decouple event boundaries related to environmental transitions from those related to object introduction (Figure 3A). Mice spontaneously move to explore novel objects, and this well-characterized behavior is used as a metric for intact memory^71^. We hypothesized that the event boundary would be defined by the object’s introduction, and therefore predicted that NE release would be related to these moments rather than the behaviors associated with individual samples of the object.

**Figure 3.**
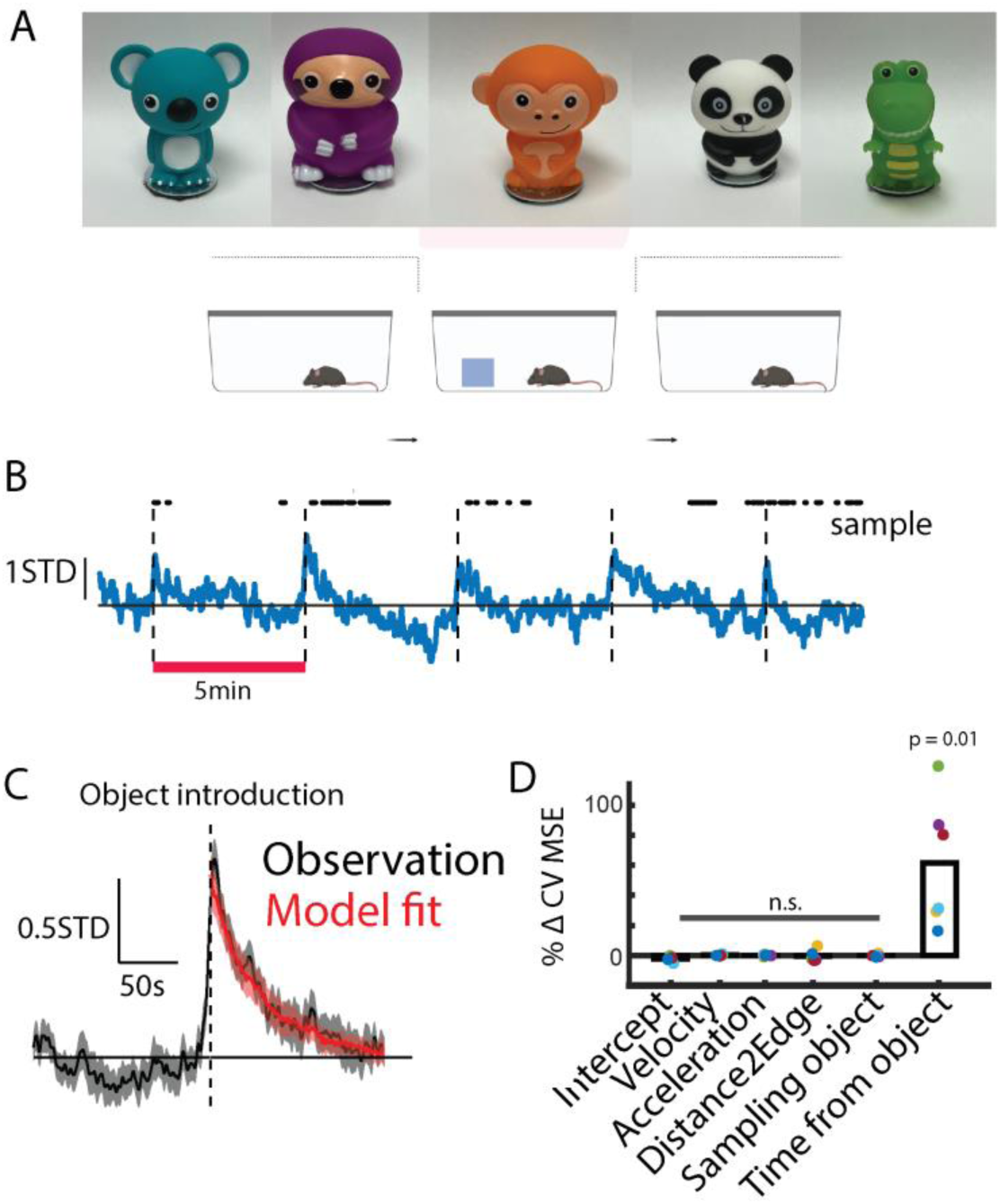
Time from object introduction controls Signal_NE_ A) Photographs of five novel objects presented to the mouse. B) Example session showing Signal_NE_ (black) aligned object introduction (dashed line) and object sampling (**·**). C) Mean Signal_NE_ measured across all object presentations (black) and cross-validated prediction from the saturated model (red). C) Change in CVMSE due to removal of various potential behavioral variables. Only removal of the terms related to time from object introduction significantly decreased model performance (*t(5) = 3.54, p =0.017*).

To address this question, a similar statistical modeling approach was adopted wherein Signal_NE_ was modeled as a function of: time from object introduction, acceleration, velocity, distance from edge of the environment, and whether or not the mouse was sampling the object. Upon introduction, each of the five objects induced a phasic release of NE (Figure 3 B,C); NE release dynamics were not systemically related to the ordinal position of the object in the sequence (Figure S3 A-C). NE release was also not coordinated with individual object samples (Figure S3D). Backward stepwise regression analysis revealed that the time from object introduction was the only term whose absence significantly decreased CVMSE (Figure 3D). These results show that changes in spatial context and introduction of salient and novel objects increase Signal_NE_, thus suggesting that NE release around both types of event boundaries may organize hippocampal neural activity.

### Novel objects do not affect Signal_NE_ around spatial context transitions

As Signal_NE_ increases around novel objects and context transitions, we next tested how the combination of these conditions affects noradrenergic signaling in the dorsal hippocampus. In addition, mice typically initiate movement to explore novelty and we sought a scenario in which mice stop to inspect something new. To achieve these goals, mice were trained to run for water on a linear track and were then presented with a novel object placed midway down the track. In these sessions, there was a baseline linear track period without novel objects, then mice were returned to the home cage and a novel object was placed on the track (Control sessions in the same subjects were run on different days without novel objects), and finally, mice were returned to the linear track. Though mice reliably stopped to inspect the novel object, no difference in Signal_NE_ was observed between the novel object and control conditions (Figure 4). Therefore, Signal_NE_ related to the familiar context transition was not affected by the presence of novel objects.

**Figure 4.**
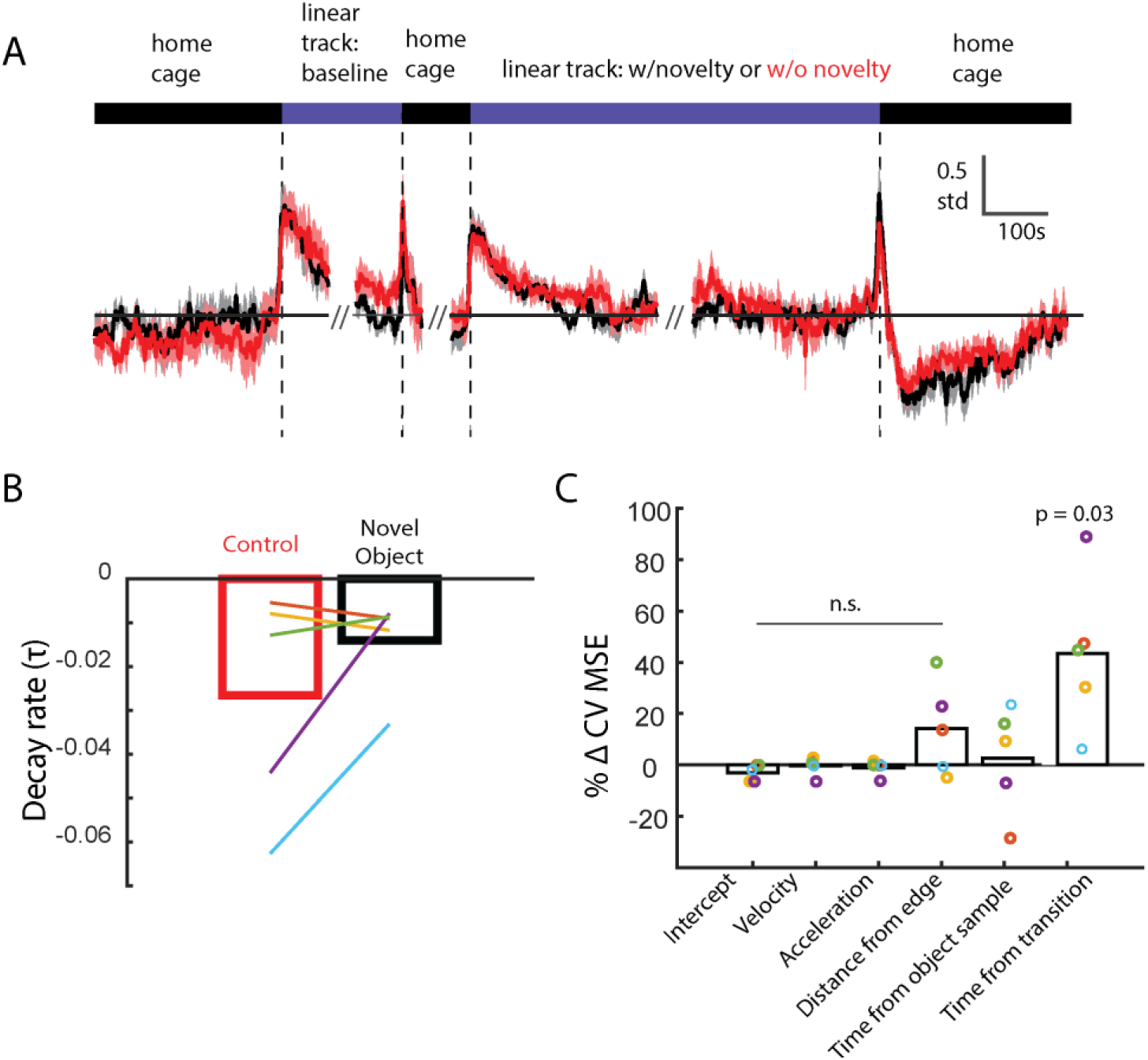
Novel objects do not affect NE dynamics after transfer to a familiar linear track. A) Mean Signal_NE_ across experimental sessions when the track was baited with a novel object (black); control sessions were run without new objects (red). B) Estimated τ describing Signal_NE_ decay after moving to the linear track did not change in the presence of a novel object (*t(4) = 1.47, p = 0.22*). C) Change in CVMSE due to removal of various potential behavioral variables. Only removal of the terms related to time from linear track transfer significantly decreased model performance (*t(5) = 3.22, p = 0.03*).

### Experience accelerates the decay of Signal_NE_ after spatial context transitions

Prior studies have found that the effect of event boundaries on the organization of memory depends on stimulus familiarity^72^ and recordings from LC neurons show rapid habituation with repeated exposures^60,62,63,73^. Therefore, we tested how the Signal_NE_ changes as a novel environment becomes increasingly familiar after repeated exposure over 10 days. Comparing the first and second days of testing, mice tended to display higher levels of acceleration, rear more often, and spend more time close to the perimeter during first-time arena exposure (Figure S4).

We adopted the same regression analysis to decouple learning-related changes in behavior from learning-related changes in NE release. As before, Signal_NE_ was estimated as a function of time from context entry, acceleration, velocity, distance from edge, and time from rearing. For each subject, for each day, we derived a point estimate of a positive β-weight associated with the gain in Signal_NE_ due to the context transition as well as a term τ that describes the rate of decay of Signal_NE_ after the event boundary. The rate of Signal_NE_ decay (τ; mixed-effects linear model, t*(73) = 2.31, p = 0.02*), and not amplitude (β; mixed-effects linear model, *t(73) = 1.16, p = 0.25*), systematically changed as a function of the number of days of experience (Figure 5 A-C). Returning the subject to their home cage was associated with an increase (β) in Signal_NE_, with a decay that was more rapid than that observed after 10 days of contextual habituation (Figure 5C). These findings show that learning alters NE signaling dynamics, either by accelerating the rate of NE clearance or by decreasing the duration in which LC neurons continue to release NE after being moved into a different space.

**Figure 5.**
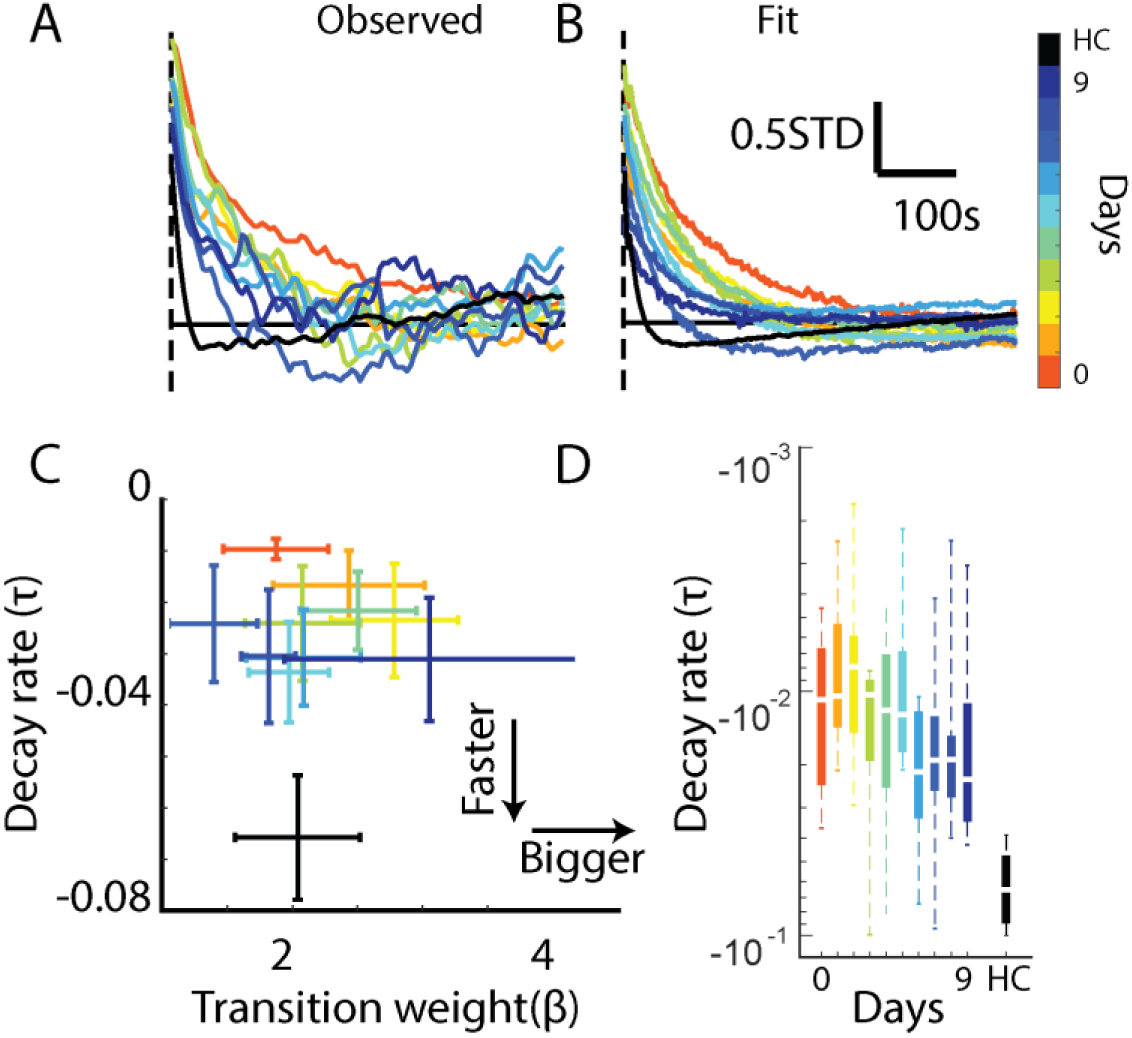
Experience accelerates Signal_NE_ decay after context transition. A) Mean Signal_NE_ plotted as a function of time from context transition (dashed line) and color coded by number of days of experience. Black trace shows Signal_NE_ recordedafter transitioning back to the home cage (HC). B) Estimated Signal_NE_ derived from the saturated model. C) Parameter estimates for the magnitude (β) and decay rate (τ) of Signal_NE_ after context transition color-coded by days of experience. D) Decay rate (τ) after transfer to the arena hastens over days of exposure (mixed-effect linear model; t*(73) = 2.31, p = 0.02*) and is most rapid during transfer to the HC (*Day N vs HC, all p ≤0.01*).

### Familiarity is not the sole determinant of the decay of Signal_NE_ after spatial context transitions

Mice were highly familiar with the linear track, yet Signal_NE_ showed a relatively slow delay. The τ_track_ was comparable to the τ_NovelEnv_ observed after 3-4 days of exposure. Moreover, there was a higher baseline Signal_NE_ maintained throughout the linear track sessions (Figure 2). We hypothesized that the dynamics of the Signal_NE_ around event transitions depend upon recent NE signaling history. To equate familiarity of the context, we compared transitions to the home cage from the linear track or the novel environments. For each session, Signal_NE_ in the homecage was modeled as a function of: context entry, acceleration, and velocity. For both linear track and novel context sessions, significant decreases in model fit were only observed after dropping the terms related to time from home cage entry (Figure S5). Signal_NE_ increased similarly around the transition to home cage after experience in the arena or linear track (Figure 6A). However, in the linear track sessions, Signal_NE_ rapidly decreased and was depressed relative to baseline for several minutes. The rate of Signal_NE_ decay was faster (Figure 6B) and the NE decrease was larger (Figure 6C) after linear track exposure as compared to experience in the arena. These results show that recent experience changes the dynamics of Signal_NE_ around event boundaries imposed by context transitions.

**Figure 6.**
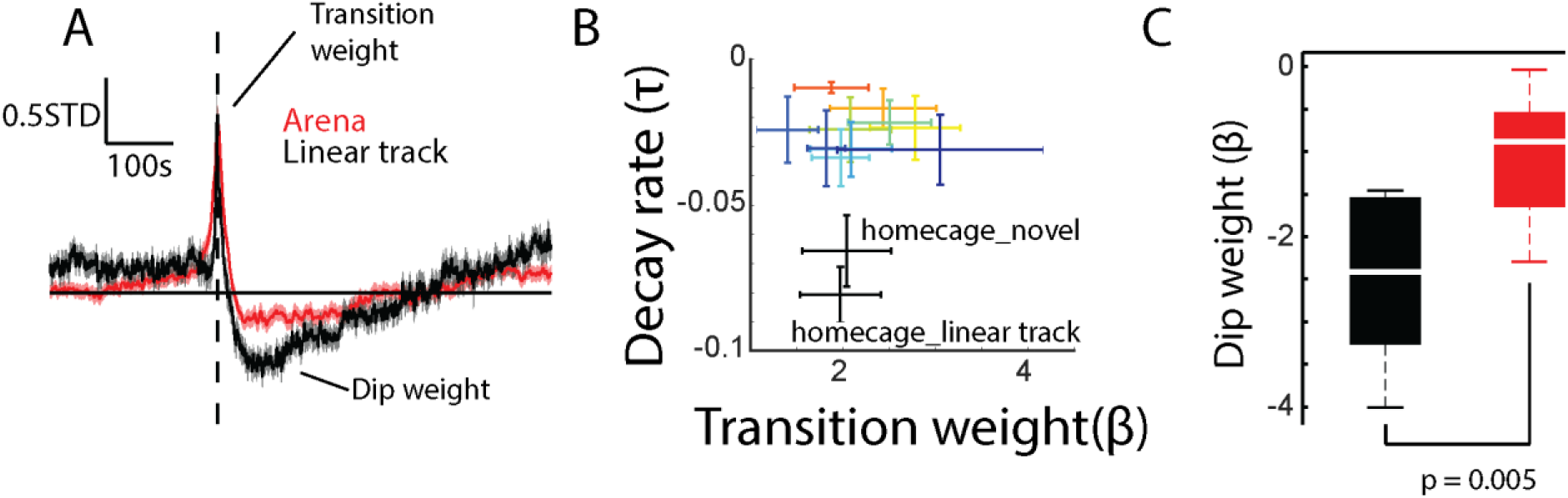
Signal_NE_ is depressed relative to baseline after periods of sustained elevation. A) Mean Signal_NE_ recorded after moving mice back to the home cage from the arena (red) or the linear track (black). B) Same data as Figure 5C with the addition of parameter estimates for the behavior of Signal_NE_ after transition to home cage from the linear track. C) The decrease in Signal_NE_ was significantly larger after transitioning mice to the home cage from the linear track as compared to from the novel arenas (*t(5) = 3.74, p = 0.005*)

### CA1 spatial code takes minutes to settle after context transition

Knowing the dynamics of NE after context transfer allowed us to search for changes in neural activity that track this time course. Modeling studies have emphasized that NE binding should increase the rate at which neural patterns change over time^27^. Using a large open-source database in which CA1 neurons were recorded as mice were transferred to novel and familiar tracks^74^, we found that in novel environments, the rate of decorrelation was indeed faster in the first minute after transfer as compared to later in the session (Figure S6 A,B). Such a relationship was not observed in a familiar space (Figure S6 C,D). Since we found strong NE release in both conditions, we doubt these changes are driven by NE.

Next, we analyzed the rate at which the spatial map settles after inducing remapping by shifting the subject from its home cage to a novel or familiar testing environment. Place fields can be observed immediately after transitioning to a new environment^75,76^, though fields can also emerge and/or change throughout experience^77^, and show other changes across repetition as well^78^. To gain an intuition for the dynamics immediately after transition, we embedded the high-dimensional population firing rate vectors (mean ensemble = 253.8 neurons, range = 191-363 neurons, bin size = 100-ms) into a 2D space. Color coding by position shows that the CA1 representational space maps the spatial layout of the environment (Figure 7A). Color coding by time shows that moments immediately following the transition are associated with unusual representations, which can be seen at the periphery of the representational state space (Figure 7B). Recognizing that single locations may have a multitude of neural representations^79,80^, we quantified the correlation of the instantaneous representation recorded at each moment relative to those recorded in the same location at any other moment throughout the session. This nearest-neighbor search revealed that early moments were associated with neural activity that poorly correlated with activity recorded in the same location later in the session (Figure 7 C,D). Representations settled into a steady state after several minutes and more rapidly in a familiar environment (Figure 7 E-G). Settling involved both an increase of activity within a neuron’s place field and a decrease in out-of-field firing (Figure S7 A,B). To ensure this representational uniqueness did not arise due to unusual behaviors during the first minutes after transfer, we calculated the absolute difference in velocity (|Δvel|_NN_) and acceleration (|Δacc|_NN_) recorded at the moments captured by the nearest-neighbor (NN) search. When comparing pairs of moments with the highest representational similarity, there was no systematic relationship between time after transfer and |Δvel|_NN_ or |Δacc|_NN_ (Figure S7 C-F). To confirm this impression, we modeled the nearest-neighbor representational similarity as a function of time from transfer, |Δvel|_NN_, and |Δacc_|NN_. Only removing time from transition significantly decreased ability to predict nearest neighbor correlations (Figure S7 E,F). Similar results were found when representational similarity was not conditioned on the mouse’s location (Figure S7 G,H). These results show that changes in representational uniqueness are more driven by time from transfer than unusual movement statistics. The time course of representational stabilization qualitatively matched that of NE decay in both novel and familiar environments suggesting a potential link between NE release and atypical spiking behavior.

**Figure 7.**
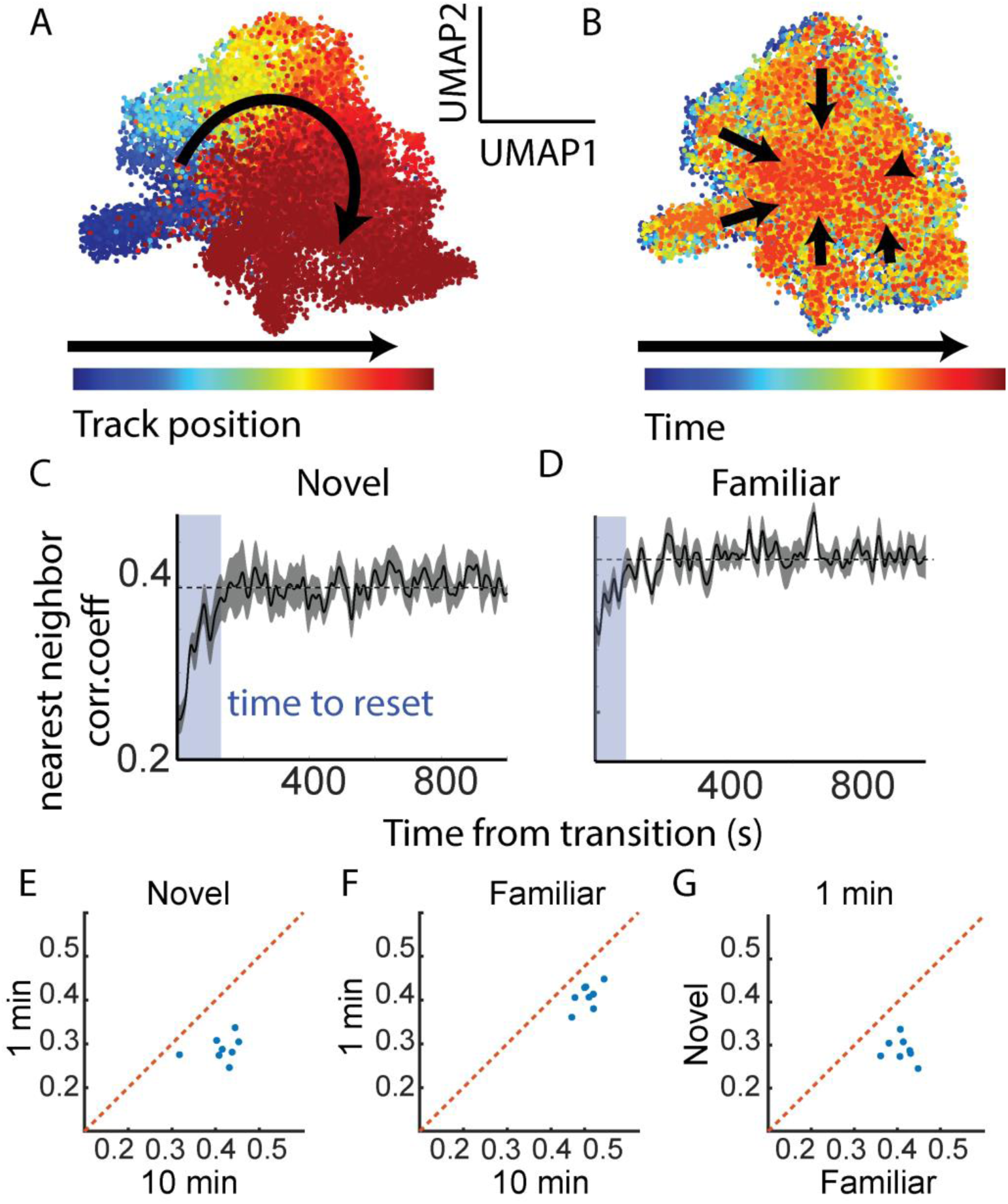
CA1 spatial code takes minutes to stabilize after context transition in novel and familiar spaces. A) Example UMAP embedding of population firing rate vectors (100-ms), color-coded by where the mouse was physically located on a linear track when the data was recorded. B) Same embedding color coded by time from context transfer. C) Representational similarity (Pearson R) of the observed population firing rate vector at each moment in a novel environment relative to the mean of the next 3 most similar vectors recorded in the same location. D) Same as Panel C recorded in a familiar environment. E) In a novel environment, the patterns recorded in the first minute were less correlated than those observed 10 minutes into the session (*t(7) = 8.05, p = 0.00009*) F) Same as Panel E recorded in a familiar environment (*t(7) = 8.20, p = 0.00008*). G) Initial representations were more correlated to their nearest neighbors in a familiar environment as compared to those recorded in a novel environment (*t(7) = 7.58, p = 0.0001*).

### No preferential reactivation of moments following transition

NE binding facilitates the induction of synaptic plasticity across hippocampal subfields^2–11^. Another body of work has shown that reactivation of waking patterns during sharp-wave ripples depends upon the same signaling pathways that mediate synaptic plasticity^43,44^, thus motivating the hypothesis that replay depends upon synaptic plasticity. Knowing when NE is likely to be present, we next asked whether the moments immediately following context transition were associated with enhanced reactivation. The population firing rate observed in each 100-ms bin was correlated with that observed during ripples before and after experience in a novel environment. These correlations were then compared to a bootstrap distribution (shuffling neural activity across ripples to break patterns of synchrony) to assess the likelihood that a particular firing rate vector would be observed more than expected if neurons fire independently of one another across ripples. Contrary to expectations, the pattern of activity observed towards the end of the session was more likely to be reactivated in the ripples that followed the experience (Figure 8 A,B). We also did not observe preferential reactivation of the moments following a transition in familiar environments (Figure 8 C,D), nor any evidence that the pattern of activity observed on the track was present in ripples recorded prior to the experience. These results suggest that enhanced NE signaling associated with context transition is not sufficient to gate entry into subsequent replay.

**Figure 8.**
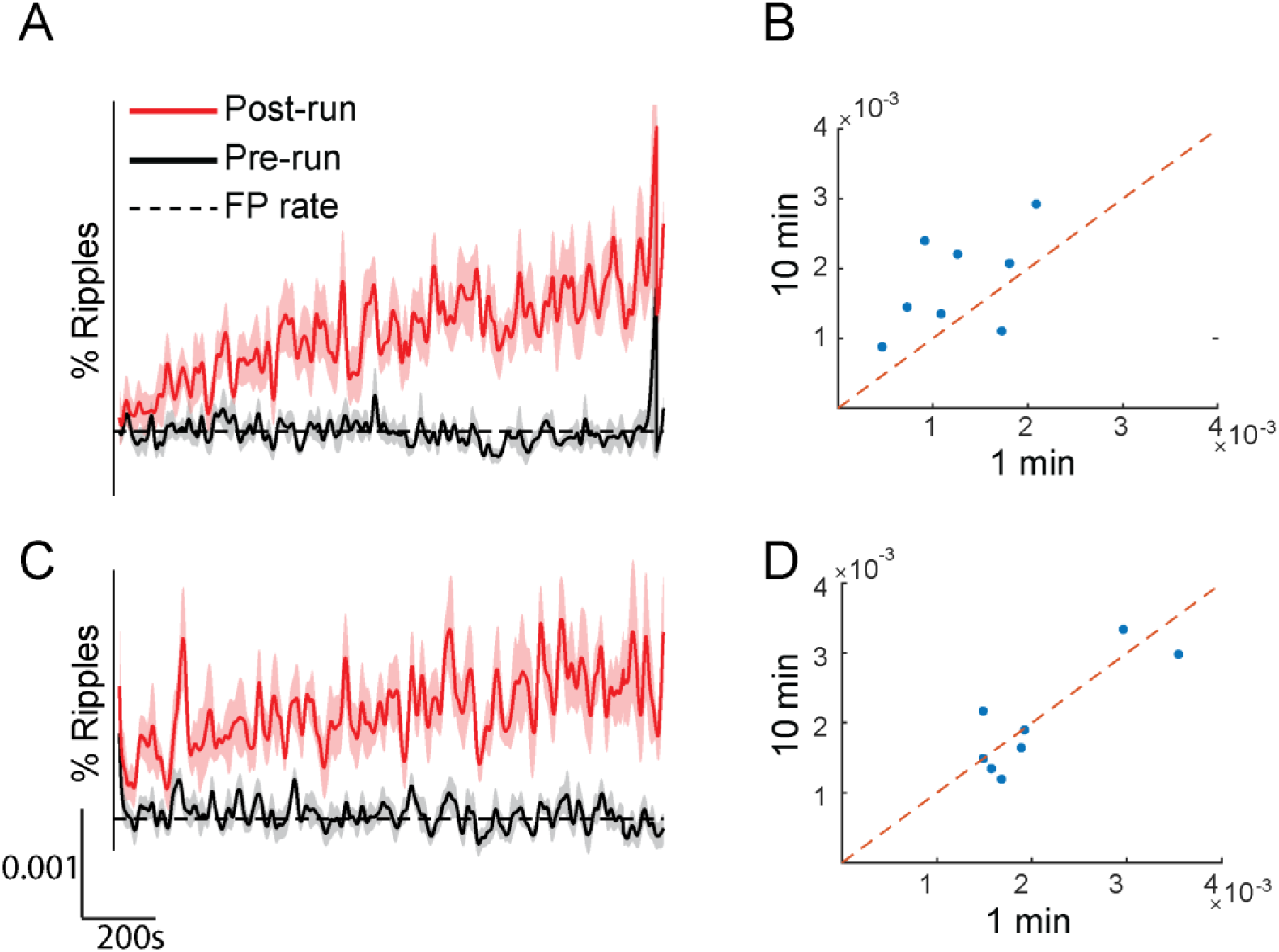
Moments immediately after transition are not preferentially replayed. A) Percentage of ripples recorded before (black) and after (red) experiencing a novel environment that showed significant reactivation of each moment after transition. Dashed line shows false positive (FP) rate. B) Moments recorded 10-11 minutes after novel context transition were more likely to be reactivated than those recorded 0-1 minutes after transition (*t(7) = 2.46, p = 0.04*). C) Same as Panel C showing reactivation rates as a function of time after transition to a familiar environment. D) There is no difference in reactivation rate for early vs late moment in a familiar environment (*t(7) = 0.40, p = 0.70*).

## Discussion

Moment-to-moment changes in extracellular NE concentration were mainly driven by the time since a salient environmental change. NE release could not be explained by fluctuations in spontaneous or conditioned mouse behavior. Familiarity accelerated the rate at which NE decayed to baseline after transitioning between contexts, while the degree of phasic NE increase at the time of transition did not systematically change with learning. In opposition to predictions from models that place a central role of NE in gating the plasticity required to alter future neural dynamics, we did not find any enhancement in the reactivation of neural patterns observed in the moments immediately following context transition, and in fact, we observed the converse – greater reactivation of the neural patterns observed later in the recording session. Analyzing the dynamics of neural coding around environmental transitions, we observed that hippocampal representations of space took several minutes to stabilize into a modal steady state. This time course was faster in a familiar environment and qualitatively mirrored that of NE release. These results support a model in which the hippocampal NE release is proportional to the deviance between the current neural representation and the steady-state attractor.

### Potential sources dictating NE dynamics

NE dynamics were well described by the sum of two exponentials, one reflecting an increase in NE release around the event boundary that decays to baseline over several minutes and another describing a decrease in NE release from baseline that recovers more slowly. This phenomenological model was able to capture complex interactions between NE release and clearance that dictate the available Signal_NE_. A temporally extended input driving NE release minutes after the event boundary is likely, since, in anesthetized preparations, the impulse response function of NE release after LC stimulation returns to baseline within tens of seconds, not minutes^81–83^. Moreover, large increases in Signal_NE_ returned to baseline quickly after transitioning to the mouse home cage. Therefore, NE clearance can occur quickly. In the awake behaving subject, however, brief optogenetic stimulation of LC produces an increase in medial prefrontal NE concentration that takes minutes to decay^53^. The mechanisms by which NE levels are maintained long after LC stimulation are unknown. The LC is the sole source of NE in the hippocampus and release dynamics are jointly dictated by changes in the firing of LC neurons and local modulation of the LC terminals. It is possible that the LC itself receives drive long after the event boundary that decreases systematically over time. Alternatively, electrotonic coupling between LC neurons may underlie phasic NE release^84^, and perhaps this electrical coupling slowly decays after event boundaries, or LC stimulation. This latter mechanism is motivated by the observation that phasic NE release is likely driven by changes in LC synchrony^85–87^. However, single unit recordings from the LC show no increase in firing rate when subjects are transferred to a familiar environment^61^, in contrast to the NE signal observed in the current study. This dissociation suggests local control of NE release independent of somatic action potentials.

In a synaptosome preparation, in which LC terminals located in the hippocampus are dissociated from LC somata, NE is released by NMDA receptor stimulation^67^, which is modulated by somatostatin^88,89^ and nicotinic^90^ receptors also located on the LC axon terminal. Somatostatin’s influence on NE release is independent of membrane depolarization^88^, thus introducing the possibility that the terminal depolarization may differ from the signal arriving to the post-synaptic neuron. Induction of synaptic plasticity can alter the levels of spill-over glutamate^91,92^ available to bind to NMDAR on LC terminals. One possible explanation for the acceleration of NE decay across days of arena exposure may relate to decreases in spill-over glutamate. If decay rates are dictated by glutamatergic stimulation of the LC terminal, future experiments should test whether these rates differ along the longitudinal axis of the hippocampus^93^. We predict slower decay in the ventral hippocampus. A diversity of decay rates (perhaps averaged in the present study) may provide more precise information about the time since an event boundary^94–96^.

We also observed significant and sustained decreases in NE release when mice were moved back to their homecage whose kinetics depended upon the recent history of the subject. NE release in the linear track was systematically elevated from baseline which likely creates decreased subsequent noradrenergic signaling resources^86^. Future studies should test whether learning after transition differs in the high and low NE states.

### NE release is enhanced around event boundaries

Event segmentation theory states that event boundaries occur at these prediction errors^97^, which coincide with an abrupt change, or reset, in ongoing activity^98^. Event boundaries have a profound influence on the organization of episodic memory. For example, memory is enhanced for the events immediately following an event boundary^99,100^. This primacy effect exists across encoding modalities^101^, and in animal studies of hippocampal-dependent spatial memory^102,103^. There are also fewer serial transitions across event boundaries during free recall^70^, which suggests segregation of memories into discretized episodes^104^. This segregation is particularly evident when networks reorganization (reset) around the transition point, as inferred by decreased correlations in multi-voxel BOLD signals^105–107^. NE released from LC terminals is known to correlate with pupil diameter^108,109^, thus providing an indirect (if imperfect^110^) assessment of LC function in people. Around event boundaries, pupils tend to dilate^1^, suggesting NE release at these times.

Direct NE measurements in animals show enhanced release in the hippocampus around conditioned and noxious stimuli, as well as following exposure to novel contexts of even handling^50–52,54,81,111^. Microdialysis studies lack the temporal resolution to dissociate whether NE release is related to specific stimuli or novelty *per se* versus the associated changes in animal behavior. Prior studies that have used GRAB_NE_ in the hippocampus have not attempted to disambiguate these possibilities.

Other recording studies have found LC neuronal activity is related to movement^55^, orienting behaviors^87^, and reward consumption^64^ and NE recordings in other brain regions have found correlations with these variables^65^. We used two techniques to isolate NE signals related to event transitions from those related to reward, movement and overall arousal. First, our statistical modeling showed across a variety of testing conditions that the time elapsed after some environmental change predicted NE release; translational movement, reward, and bouts of exploratory behavior (rearing or object exploration) were poor predictors of Signal_NE_. Next, we developed different protocols in which exploratory behaviors either involved the initiation or the interruption of movement. In neither case did we observe time-locked NE release around bouts of exploration.

Arousal or attention also seem to be unsatisfactory explanatory cognitive constructs to explain the dynamics of hippocampal NE release observed in the present study. In as much as these mental states can exist or be measured in the rodent, situations in which mice systematically engage in more exploration did not change the time course of NE decay after context transition (Figure 4). Instead, in all cases tested, the hippocampus NE release corresponded to the time elapsed from an unpredicted salient environmental change (context shift or object introduction). Notwithstanding, in a subset of subjects, we did observe transient changes in NE release around rearing events. Though this was not significant at the group level, we speculate that the degree of NE release may be related to the nature of the information acquired during the environmental sampling.

### The LC contribution to long-term changes in neural coding

The LC influences memory formation through the co-release of dopamine^61,69,111–115^ and NE^2,54,116–119^. The modulation of late-phase synaptic plasticity, e.g. through synaptic tag-and-capture mechanisms^3,120^, has long been emphasized as the dominant role by which catecholamines may gate entry of new information into long-term memory^4,9,11,20,61,112,113,121–124^. SPW-R replay is a prominent electrophysiological correlate of experience that depends on plasticity-related processes^43,44^. Since stimulation of the midbrain enhances replay^47^, we hypothesized that NE may also enhance future reactivation. This prediction was not correct, as we did not find any evidence that the neural activity observed following context transition was preferentially reactivated. In fact, we saw that *later* moments were more likely to be reactivated in post-RUN ripples. This reactivation bias is likely due to the autocorrelation of the brain over time in which the neurons active at any given time are more likely to continue to be active due to consistencies in the external environment (or internal milieu) and the slow turn-over in proteins that affect intrinsic excitability^125–127^. Though we did not quantify replay of the temporal sequences of cell assemblies, a prior report using the same data also failed to observe enhanced replay of moments following transition^128^. Therefore, if a primacy effect occurs after context transitions, it is unlikely to be mediated by, or reflected in, enhanced replay of these moments. Others have found that LC stimulation promotes place field accumulation, but only in the present of natural reward^69^. NE is therefore likely to act in concert with other signals to promote long-term changes in neural coding during exploration and during ripples.

### Changes in neural coding around event boundaries

We observed that immediately after an environmental transition, the spatial representation was relatively unique, and settled into a steady-state spatial code over the course of minutes. In familiar spaces, the neural patterns observed in the early moments were more similar to the ultimate steady state. When subjects move between environments, hippocampal place fields remap which involves changes in which neurons express place fields, alterations in which subsets of neurons fire together, and reorganization in the distances between the place fields of simultaneously recorded neurons^33,39^. This remapping can occur rapidly, with the reset signal driven either externally – when stimuli signal changes in how subjects should behave within the space^36,129^ – or internally when multiple reference frames must be simultaneously maintained^130,131^. During such rapid remapping competing ensembles “flicker” before settling into a steady state^129,132^. Manually moving subjects between environments also induces remapping^68^. Place fields may be observed on the first trial in a novel environment^75,76^, but previous studies have found that extended exposure modifies the hippocampal representation of space in several ways: new fields may be added^77,133^, field asymmetry changes^78^, and firing reliability is enhanced^76^. Other changes may occur in the presence of appetitive^34^ or aversive stimuli^35^.

The time course for reset around transition, in which neural activity reached its steady state, qualitatively matched that of NE release. It is possible that NE perturbed neural activity away from the stored attractor. The seminal work of Segal and Bloom showed that electrical LC stimulation acutely silenced most hippocampal neurons^24,25^ while enhancing the firing rate of those neurons that fire in response to various stimuli. In anesthetized rats, LC activation causes an increase in the excitability of CA1^23,134^ and dentate gyrus^20^ neurons, as measured by the amplitude of the population spike after afferent bundle stimulation. *Ex vivo* low-frequency optogenetic stimulation of LC terminals likewise causes an increase in CA1 intrinsic excitability^23^. These acute effects are all blocked by β-adrenergic receptor antagonists. Therefore, NE-related changes in gain/excitability may cause deviations from a stored neural representation.

Alternatively, prominent models stipulate that area CA1 could be key in the generation of a memory-related surprise signal that redirects attention and drives the release of neuromodulators^4^. In these models, an error signal originates from a “comparator” structure in CA1^4,135,136^. This hypothesis was motivated by the observation that CA1 neurons are activated by contextual novelty^137^, novel objects^138^, and novel configurations of familiar objects^139^. Unexpected violations of a learned sequence also cause robust activation of CA1 neurons^140^, an output that may be used to signal prediction error to arousal circuits^141,142^. This error signal may drive NE release through local modulation of LC terminals, or through polysynaptic pathways (e.g. via the paraventricular hypothalamus^143,144^). We speculate that an error signal should be proportional to the difference between the instantaneous and steady-state neural representation.

We observed relatively unique neural patterns immediately following event boundaries. Computational models predict that “pattern separation” yields enhanced memory by virtue of creating neural traces that are less susceptible to interference^145^. The hippocampal activity patterns observed soon after the transition provide a neural timestamp for those moments that may, in turn, underlie the enhanced subsequent recall that defines the primacy effect.

### Limitations

The main limitation of the present study is that NE and neural coding were not studied in the same subjects. Future studies should combine recording modalities and causally link the changes in neural activity and NE signaling through perturbation studies that up- and down-regulate NE and test for changes in hippocampal coding through the lens of representational uniqueness.

Another important limitation of the present study is the lack of *in vivo* calibration of the GRAB_NE_ sensor. First, all measurements here are relative to baseline. Future studies should estimate how emission intensities scale with NE concentration *in vivo*. Relatedly, the sensor is expressed everywhere on the neuron, thus providing a read-out of a signal that may not actually be available to the post-synaptic cells. Though most NE signaling occurs via “volume transmission”, noradrenergic receptors do show laminar specificity^146^ that is not honored by the membrane insertion patterns of the GRAB_NE_ sensor. Finally, the sensor has fast onset (τ_on_ = 0.09 s) and slow offset kinetics (τ_off_ = 1.93 s)^53^. Additionally, we smoothed the Signal_NE_ which, combined with limitations of the sensors, impose some limitations on the rate of behavioral fluctuations that may be captured in our analyses. The temporal resolution of the sensor has not been calibrated against amperometry or fast cyclic voltammetry, but once such experiments have been done, a deconvolution kernel may be developed to correct for binding kinetics.

Finally, the results have implications for a larger literature focusing on memory enhancement for the events that occur after an event boundary. We define a minutes-long time window in which a potential noradrenergic-dependent primacy effect may be expected, however, we did not quantify learning gains as a function of time from an event boundary. Relating the present observations to memory is an important future direction.

### Conclusion

We found that the primary driver of NE release in the dorsal hippocampus is time from some salient environmental change. When NE is elevated, neural activity differs from its steady state, which may promote subsequent retrieval of these moments associated with relatively unique neural representations. Event segmentation disturbances have been observed in a variety of disorders, including: ADHD^147^, schizophrenia^148^, and Alzheimer’s Disease^149^ (a disease in which the LC is particularly affected^150,151^); as well as in normal cognitive decline in aging^149^. Trauma can also affect noradrenergic signaling in the hippocampus^152^, which affects how we respond to and cope with stress^153^. Future studies that relate NE release to hippocampal network remapping/reset will provide important insight into the comorbid attention and memory deficits associated with these disorders.

## Methods

### Fiber photometry

#### Subjects

C57BL/6J mice (N = 8 mice, N = 3 female) were implanted at 3-6 months-old. Data was acquired for up to a year after implantation with no change in signal quality across this extensive timeline. Two surgeries were performed at least two weeks apart, the first to deliver the GRAB_NE_ sensor via AAV infusion and the second to implant a fiber optic stub. After viral injection, animals were housed individually on a regular 12:12 h light:dark schedule and tested during the light cycle. Following one week of recovery from the second surgery, mice were recorded at most 5 days/week for up to a year before being euthanized with a sodium pentobarbital cocktail (FatalPlus®, 300 mg/kg I.P.) and transcardially perfused with 4% paraformaldehyde. All experimental procedures were performed in accordance with the National Institutes of Health Guide for Care and Use of Laboratory Animals and were approved by the University of New Mexico Health Sciences Center Institutional Animal Care and Use Committee.

#### Viral injections and fiber implant

Mice were deeply anesthetized with isoflurane (1.5-2%in pure oxygen) and GRAB_NE_ was delivered by injecting AAV9-hSyn-NE2h (titer: ≥ 5×10¹² vg/mL, WZ Biosciences, MD USA)^53^ unilaterally into the left dorsal hippocampus. Two coordinates were used, both with reference from bregma: coordinate 1 (N = 2 mice) A/P: -2.3, M/L: -2.0 D/V: -1.4 and -1.2 from the brain surface; coordinate 2 (N = 6 mice) A/P: -2.0, M/L: -1.5 D/V: -1.3 and - 1.1 from the brain surface. Coordinate 1 yielded higher signal-to-noise; signals recorded from both coordinates showed the same qualitative dynamics around event boundaries. In all cases, the virus was injected at two depths each at a volume of 150-nL and a rate of 30 nL/min using a Nanoliter 2020 Injector (WPI). At least two weeks later, fiber optic stubs (10 mm borosilicate mono fiber-optic cannulas from Doric lenses; MFC_400/430-0.66_10.0mm_MF1.25_FLT) were implanted at the injection site. To secure the stubs to the subject, the surface of the exposed skull was covered with C&B Metabond® (Parkell, NY USA), and the sides of the exposed fiber-optic cannula were coated in Unifast LC dental acrylic (SourceOne Dental, Inc, AZ USA) for stability. Finally, clips (Neuralynx, AZ USA) were added to minimize motion artifact due to slippage at the mating sleeve. Postoperatively, animals received a single injection of 0.1-mg/kg of buprenorphine (S.C.) and again as needed for the next 1-3 days.

#### Fiber photometry recording procedures

Prior to the first recording session, we allowed a minimum of three weeks from the viral injection procedure to allow the virus sufficient time to transfect and express. Signals were captured with a LUX RZ10X processor running the Synapse software (Tucker-Davis Technologies, FL). Experimental (465 nm, carrier frequency = 330 Hz) and isosbestic (405 nm, carrier frequency = 210 Hz) wavelengths were combined using a fluorescent MiniCube (FMC4_IE(400-410)_E(460-490)_F(500-550)_S; Doric, QC Canada) and delivered to the subject with a 4-m low auto fluorescence mono fiber-optic patch cord (core = 400-µm; NA = 0.57; Doric, QC Canada). Excitation intensity of the isosbestic and experimental wavelengths was adjusted to equalize emission intensity, which was sampled at 1017.3 Hz.

#### Behavioral procedures

##### Novel arena

On the first day, mice were transferred to three novel arenas (dimensions in Figure S1). First, a 10-minute homecage (HC) baseline was captured, then mice were manually transferred to a novel arena (Context A) and back to their homecage for 10 minutes. This procedure was performed again for Contexts B and C (HC-Context A-HC-Context B-HC-Context C-HC). On following days, a 10-minute baseline period was run, followed by 10 minutes of exposure to Context A, and another 10 minutes in the home cage (HC-Context A-HC). On Day 10, the procedure from the first day was repeated.

##### Spontaneous Object Recognition

On Day 0, mice were allowed to acclimate to a clean and empty cage for 30 minutes. This cage had a hook-and-loop fastener for later object placement. On Day 1, we recorded a 10-minute baseline in the clean and empty cage. Then, five novel objects were sequentially affixed to the hook-and-loop fastener in the cage, each for five minutes with no interval between objects. After the fifth object was removed, the animal remained in the empty cage for another 10 minutes.

##### Linear track

Water-restricted mice were trained to run laps on a 1.2m linear track for water reward (15µL) which was delivered at each end of the track after mice crossed an IR sensor to trigger a wall-mounted solenoid. Mice ran between 3-17 laps (mean = 8.1 laps) in 286-1500s (mean = 658s). In these sessions (N = 110), there was a 10-minute homecage period before mice were transferred to the linear track. Once mice stopped running for water for at least 30s, they were returned to the home cage for 10 minutes. Following data acquisition, mice were given *ad libitum* access to water in their home cage for 15 minutes and weighed to ensure no more than 15% loss of baseline body weight.

#### Drug infusions

Desipramine hydrochloride (Bio-Techne Corporation, MN USA) was injected (I.P.) at a concentration of 10mg/kg (1 mg/ml) in normal saline (0.9%). Yohimbine hydrochloride (Sigma Aldrich, MO USA) was injected (I.P.) at a concentration of 4-mg/kg (0.4 mg/ml) in normal saline. For recordings with drug injections, a 10-minute baseline was captured before injections with either drug or vehicle.

#### Signal Analysis

##### Estimation of Signal_NE_

The demuxed experimental and isosbestic signals both exhibited evidence of photobleaching, though with different decay rates. Therefore, we fit a double exponential to the first 10 minutes of each signal to estimate and extrapolate a mean signal which was subtracted from the observed emission intensities. Next, the isosbestic was scaled to the experimental signal using standard linear regression. The isosbestic was then subtracted from the experimental signal, and the mean and standard deviation were calculated over the first 10 minutes. These values were used to normalize Signal_NE_ which is measured in terms of baseline standard deviations from the baseline mean. Finally, the signal was smoothed with a Gaussian kernel (1-s s.t.d.).

We opted against a sliding window dF/F calculation, as we did not want to impose a minutes-long timescale to our analysis and we opted against divisive normalization directly to the isosbestic as photobleaching dominated the fluctuations in the isosbestic signal and this rate differed from that experimental signal^154^. We adopted the mean and standard deviation from the baseline period (rather than the entire session), as some of our experimental conditions (e.g. desipramine infusions) dramatically changed the mean Signal_NE_ values over long periods of time. We are aware that subtractive isosbestic correction (instead of divisive) may distort the relative amplitudes of signals recorded early versus late into the session^155^. These concerns are mitigated here as the main decreases in emission intensity due to photobleaching occurred within the 10-minute baseline period. Moreover, we observed stable responses across ∼1-hr of recording (e.g. see Figure 1C) and a reliable return to baseline Signal_NE_ values in the final home cage recordings.

##### Statistical modeling of Signal_NE_

Signal_NE_ at each moment was estimated as a function of various behavioral variables which differed according to the testing paradigm.

In the novel arena experiments, Signal_NE_ was estimated as a function of acceleration (*acc*), velocity (*vel*), normalized distance from the edge (*distedg*), time from context transfer (*t1*), and time from rearing onset (*t2*), see *Equation 1*. Acceleration and velocity were calculated using a second-order Kalman filter of the head location (right and left ear locations estimated with DeepLabCut^156^). Normalized distance to the edge was calculated as the distance to the nearest edge divided by the maximum distance to an edge possible. In some cases, the animal could extend its head beyond the wall of the arena and these values were coded as negative.

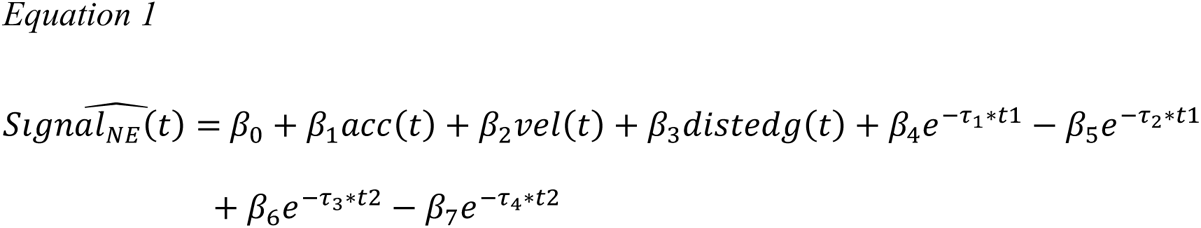

Time from transition/rearing was modeled with two terms: a positive term β_4/6_ with a fast exponential decay τ_1/3_ and a negative term β_5/7_ with a slower exponential decay τ_2/4_. To avoid degeneracy, τ_1/3_ was bounded between 0.1-0.001 and τ_2/4_ was bounded between 0.001-0.0001. All β values were bound at ±10 s.t.d. Point estimates for the 12 free parameters (β_0-7_ and τ_1-4_) were calculated with MATLAB R2021b using the fmincon non-linear optimizer against a regularized objective (Equation 2) defined by the mean squared error (MSE) with a penalty for model complexity (λ = 0.001). Fits were robust to initial conditions.

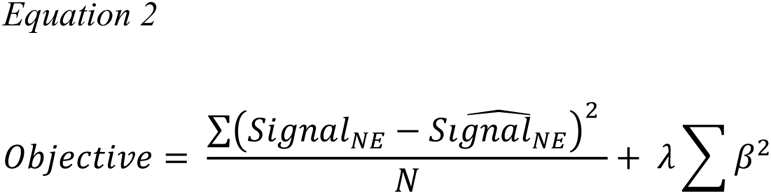

We performed 50/50 cross-validation, with the model trained on even days and tested on odd, or vice-versa. The cross-validate mean-squared error (CVMSE) was used to assess model fit (the regularization term is dropped here).

To assess the importance of each behavioral independent variable (and intercept), we excluded all terms related to those variables in a backwards stepwise regression analysis. For example, removing time from context transfer removed four terms: β_4_, β_5_,τ_1_,τ_2_. The cross-validation employed here ensures that model performance should not suffer more simply due to removing more free parameters, as demonstrated by the stability of the model after removing the four terms related to rearing (or reward in the case of the linear track). CVMSE for the saturated and reduced model was compared by computing the percent change in CVMSE.

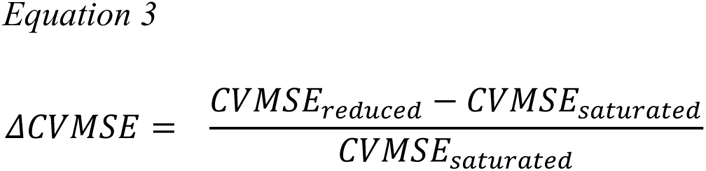

A similar approach was adopted for modeling Signal_NE_ during novel object exposure, except we included a binary indicator function for whether the mouse was sampling the object (snout touching the object) and the time from event boundary, *t3*, was the time from object introduction; we dropped the term related to rearing. Parameters were estimated for each subject and 50/50 cross-validation was done by splitting each session in half (first half training, second half test).

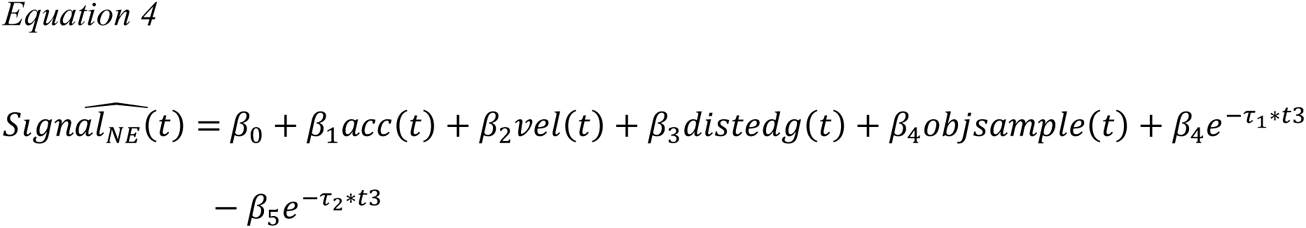

For the linear track, we considered: velocity, acceleration, distance from edge, time from transfer to the track (*t1*), and time from reward (*t4*). Parameters were estimated for each subject and cross-validation was done by considering even training and odd testing days (or vice versa).

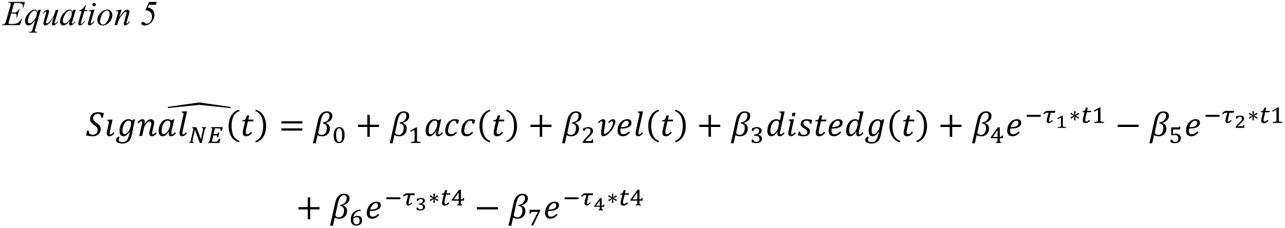

In all cases, to determine the significance of a parameter’s removal, we performed Student t-test on the CVMSE values (testing against *h_0_* CVMSE = 0) with degrees of freedom defined by the number of subjects. To compare changes in parameters across days, we used a mixed-effects linear model, with days of exposure defined as a fixed effect and subject as a random effect. We modeled the relationship with random slopes and intercepts.

### Electrophysiology

#### Electrophysiology subjects

Data was downloaded from The Buzsaki Lab Databank (Project: Place field-memory field unity of hippocampal neurons)^157^. As described in Huszar et al.^74^, chronic recordings were performed from freely moving adult C57BL/6J mice (N = 3 mice; subjID: e13_26m1, e15_13f1, e16_3m2) using high-density ASSY Int64-P32-1D or ASSY Int128-P64-1D silicon probes (Diagnostic Biochips, MD USA). In these experiments, probes were implanted over the right dorsal hippocampus (A/P -2.0, M/L +1.7) and lowered to the deep neocortical layers, while the drive was cemented to the skull. A stainless-steel screw was placed over the cerebellum for grounding and reference. Neural signals were recorded in the homecage while probes were lowered into the CA1 pyramidal layer, which was identified physiologically via the sharp wave polarity reversal. Neural data were amplified and digitized at 30-kHz using Intan amplifier boards (RHD2132/RHD2000, Evaluation System, Intan Technologies, CA USA). The complete dataset is available at https://dandiarchive.org/dandiset/000552/0.230630.2304. All experiments were conducted in accordance with the Institutional Animal Care and Use Committee of New York University Medical Center (IA15-01466).

##### Behavioral testing

Over weeks, mice were over-trained on a spatial alternation task in a figure-eight maze (see Huszar et al. 2022, for full details). Animals were water restricted before the start of experiments and familiarized with the figure-eight maze. Mice were trained to visit alternate arms between trials to receive a water reward in the first corner reached after making a correct left/right turn after which, a 5-s delay in the start area was introduced between trials. To explore the reorganization of place tuning across different environments, the same mice were introduced to novel environments after running in the familiar figure-8 maze. In the sessions analyzed here (N=8), animals underwent recording sessions consisting of a ∼120-min home cage period, running on the familiar figure-eight maze, ∼60-min home cage period, running in a novel environment, followed by a final ∼120-min home cage period. In some sessions, animals were exposed to two distinct novel environments, with a ∼60-min home cage period in between (only one transition to a novel environment was chosen per session to analyze here). We considered transitions to novel linear tracks (N = 3 sessions), novel figure-8 mazes (N = 3 sessions), and a novel arena (N = 1 session). Mazes were placed in distinct recording rooms, or in different corners of the same recording room, with distinct enclosures to ensure unique visual cues. Mouse position was captured with head-mounted red LEDs.

##### Spiking analysis

Spikes were extracted and classified into putative single units using KiloSort1^158^ and manually curated in phy^159^. Pyramidal neurons were separated from interneurons based on waveform shape and bursting statistics and only pyramidal cell spiking was analyzed.

#### ACG slope analysis

Population firing rates were calculated in 100-ms bins by counting the number of spikes observed in that period and then z-scoring over the first 1000-s after transfer. All vectors within a session were correlated with one another to generate a similarity matrix of Pearson R correlation values. We considered the drop-off in population firing rates vector correlation over a 10-s period using a 100-s moving average with an exponent with three free parameters (β, τ, c).

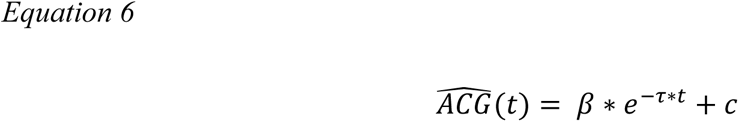

#### Reset analysis

At each 100-ms moment, we asked where was the subject in space, and what were the 3 most similar population firing rate vectors – as assessed from the similarity matrix of Pearson R values – recorded in that location (minimum occupancy = 1-s). The mean of this nearest-neighbor (NN) search was saved as the measure of representational similarity of that moment to all others, conditioned on the location of the mouse and smoothed with a 1-s Gaussian kernel.

To control for movement, we additionally calculated the mean absolute difference in velocity (|Δvel|_NN_) and acceleration (|Δacc|_NN_) for the time bins with the highest population firing rate vector correlations, i.e. those identified by the nearest-neighbor search above. If low correlations in our NN search were driven by unusual movements, we would anticipate this to be reflected by large deviations in |Δvel|_NN_, and |Δacc|_NN_. Therefore, we estimated the NN correlation as a function of time from transition, |Δvel|_NN_, and |Δacc|_NN_.

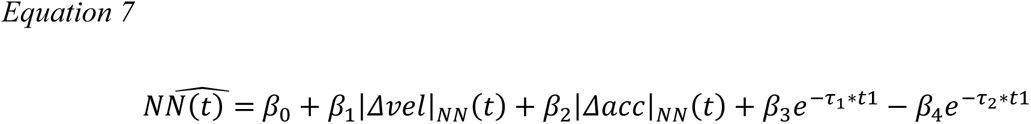

Cross-validation was done by withholding each session from the training dataset and reporting the CVMSE for each withheld session.

#### Place field detection

Mouse location was binned in 1×1 cm bins and the mean normalized firing of each neuron (as described above) was calculated in each location. During moments when velocity exceeded 5 cm/s, the mean normalized firing rate was calculate for each bin with more that 1-s occupancy. Place field bounds were defined as regions with > 5 Hz firing rate (i.e. using an unnormalized firing rate threshold).

#### Ripple detection

Broadband LFP was bandpass filtered between 130 and 200 Hz using a fourth-order Chebyshev filter, and the normalized squared signal was calculated. SPW-R maxima were detected by thresholding the normalized squared signal at 5 s.t.d. above the mean, and the surrounding SPW-R start and stop times were identified as crossings of 2 s.d.t. around this peak. SPW-R duration limits were set to be between 20 and 200 ms. See Huzsar et al.,^74^ for full details.

#### Reactivation analysis

For each ripple recorded within 30 minutes of the beginning of the session and within 30 minutes after the session, a population firing rate vector was calculated by summing the total number of spikes emitted from each unit and dividing by the duration of the ripple. Next, these population firing rate vectors were correlated with those recorded on the track (in 100-ms bins). To assess whether the observed Pearson R was greater than expected by chance, a bootstrap null distribution was created by shifting each neuron’s activity observed on any given ripple to a random other ripple observed during the session, thus preserving the single-cell mean ripple recruitment rate, but destroying patterns of synchrony observed across the ensemble. This procedure was repeated 1000 times, so that we could ask, for each ripple, if the observed Pearson R greater than 99.9% of the shuffles. We report the percentage of ripples in which each moment shows significant reactivation before and after experience with a false positive rate = 0.001.

## Acknowledgments

This work was funded by NIMH R00MH118423. I.C. was supported by IU Hutton Honors College. We are grateful for discussions with Marc Howard and Horacio Rotstein throughout the preparation of this manuscript. We are grateful to Roman Huszár for proving the electrophysiological data and useful comments on the manuscript.

## Supplemental Figures

**Figure S1.**
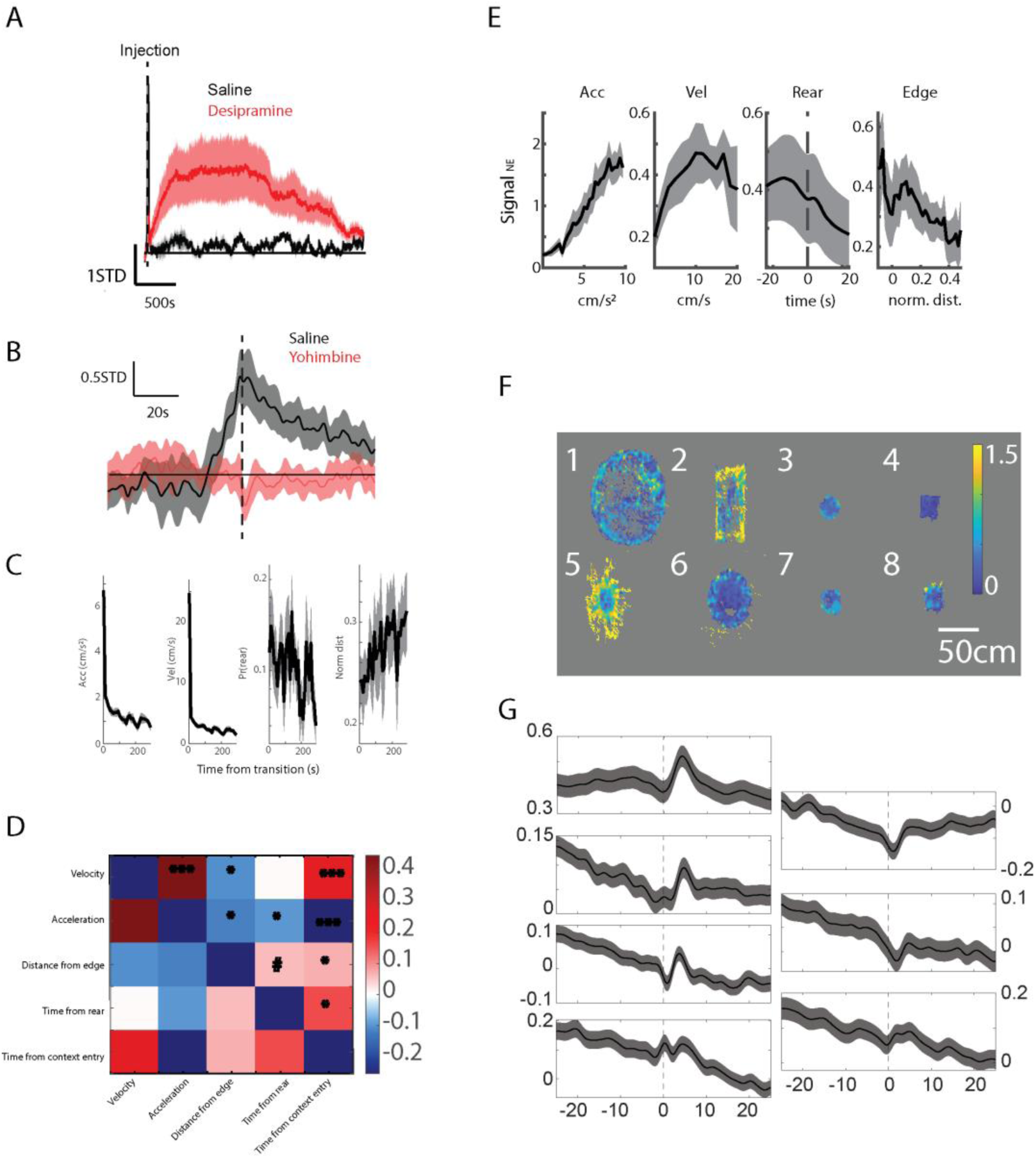
Validation of GRAB_NE_ sensor and behavioral correlates of Signal_NE._A) Signal_NE_ increases after injection of desipramine. B) The normal increase in Signal_NE_ after context transition is eliminated after injection with yohimbine. C) Fluctuations in behavior as a function of time after context transition. D) Time series correlations (Pearson R) in independent behavioral variables used to predict Signal_NE_. E) Signal_NE_ plotted as a function of different behavioral variables. F) Signal_NE_ plotted as a function of mouse position in each of the novel arenas. G) Signal_NE_ plotted for each mouse as a function of time around rearing (data for one subject was not available).

**Figure S2.**
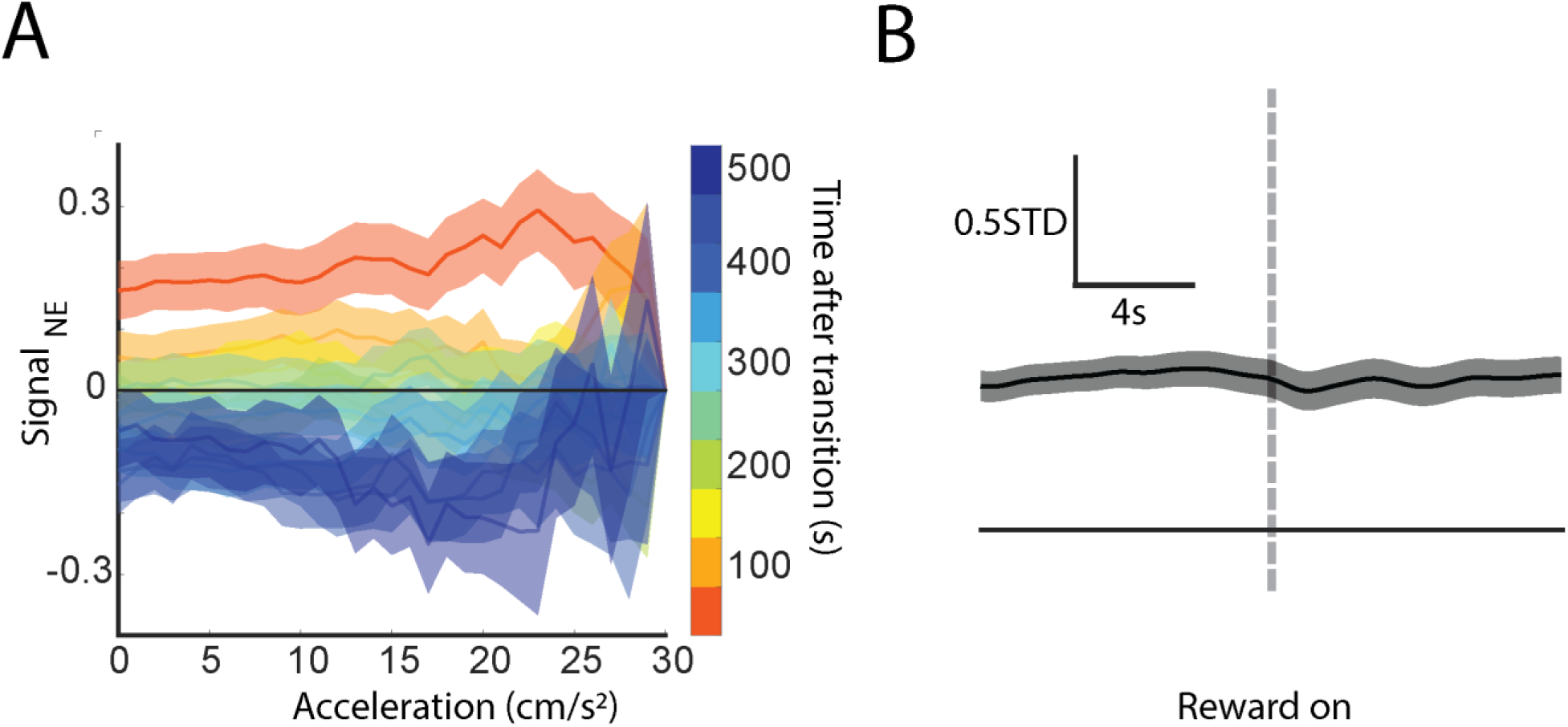
No change in Signal_NE_ due to acceleration nor reward delivery on a linear track. A) Mean Signal_NE_ plotted as a function of acceleration conditioned on time after transition. B) No change in Signal_NE_ after reward delivery (dashed line).

**Figure S3.**
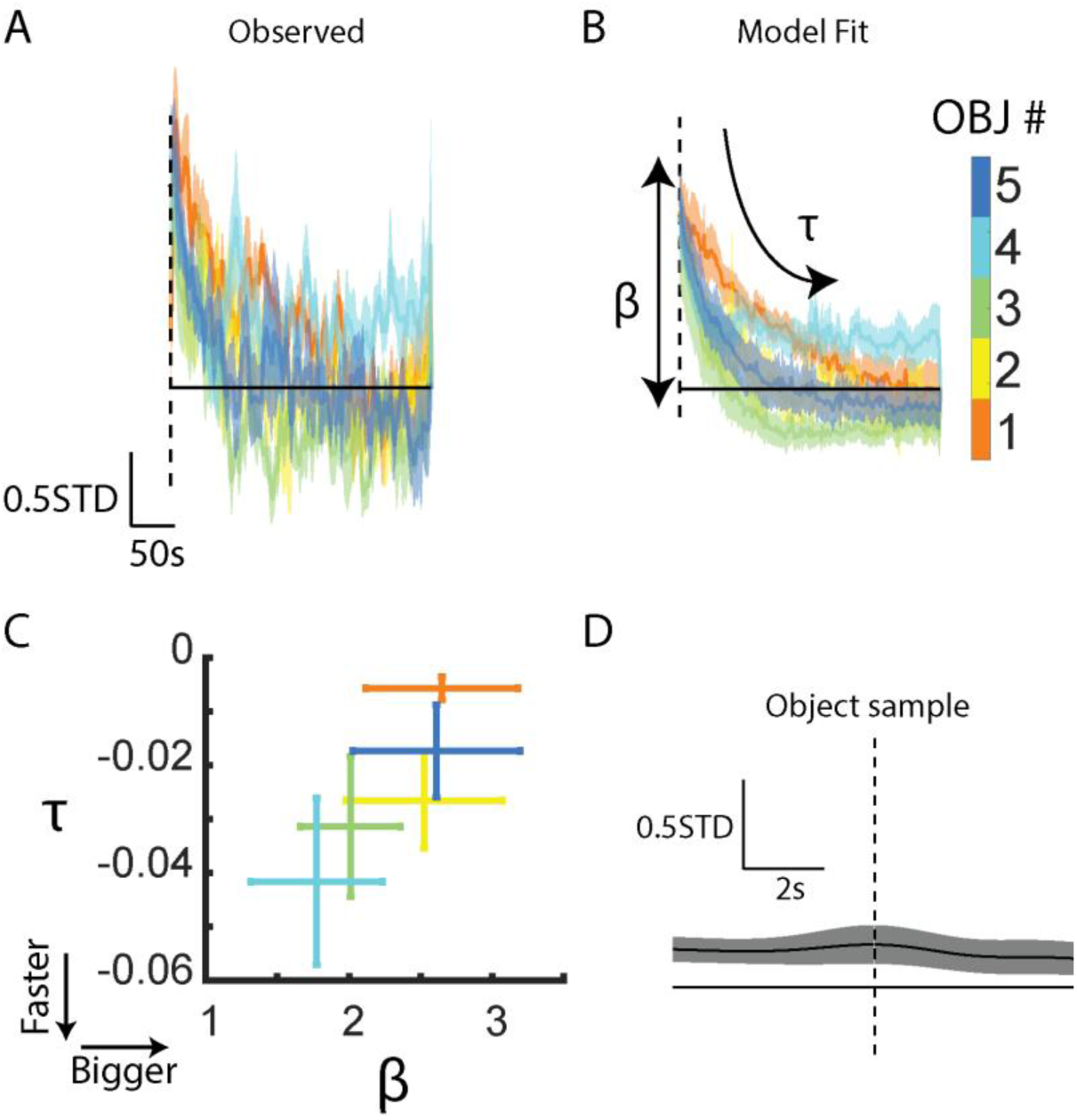
Signal_NE_ is related to object introduction, not sampling. A) Observed mean Signal_NE_ around each object’s introduction. B) Estimated fits derived from the saturated model. C) Mean ± SEM point estimates for the increase (β) and decay (τ) in Signal_NE_ around introduction of each object. D) Mean observed Signal_NE_ around each object sample.

**Figure S4.**
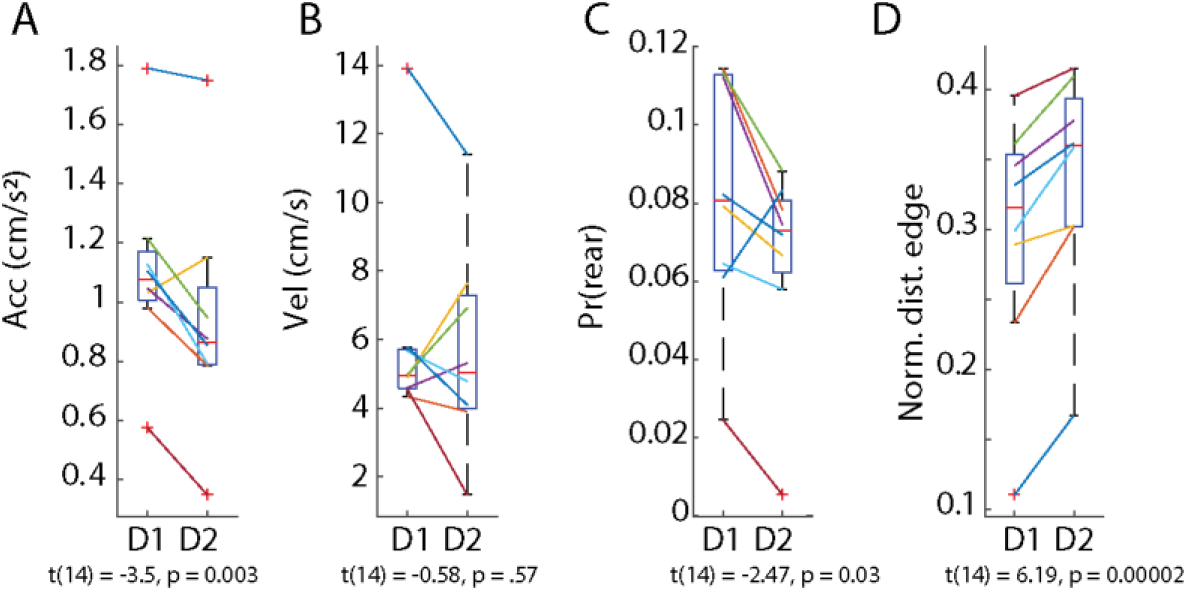
Change in behavior across days. Change in A) acceleration, B) velocity, C) propensity to rear, and D) distance to the edge across day 1 (D1) and day 2 (2) in the novel arena.

**Figure S5.**
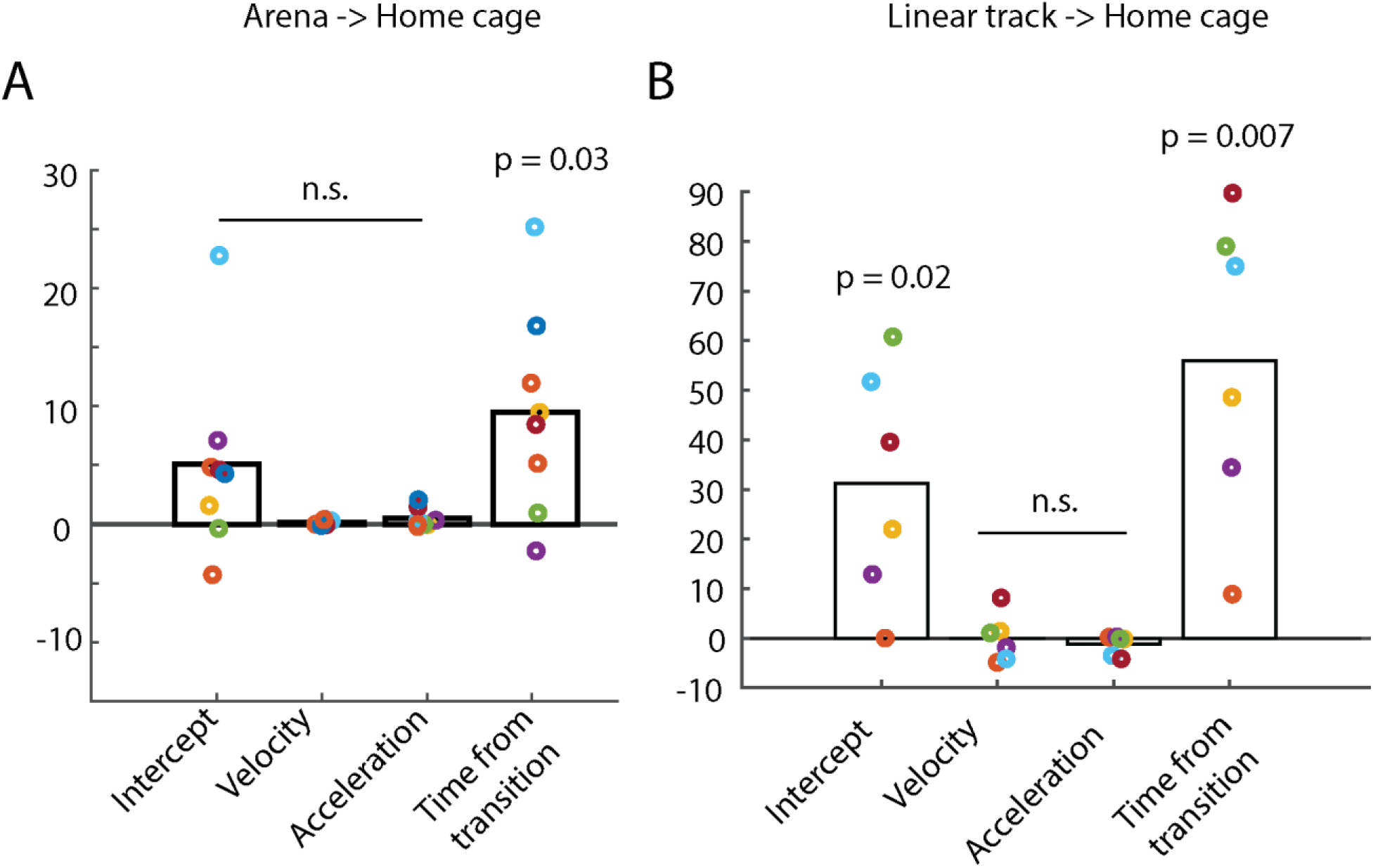
Time from context transfer explains Signal_NE_ in the home cage. A) Change in CVMSE due to removal of various potential behavioral variables. Only removal of the terms related to time from home cage track transfer from the arena significantly decreased model performance, (*t(7) = 2.62, p = 0.03*) B) Same as Panel A with transitions to the home cage from the linear track (*t(5) = 4.44, p = 0.007*)

**Figure S6.**
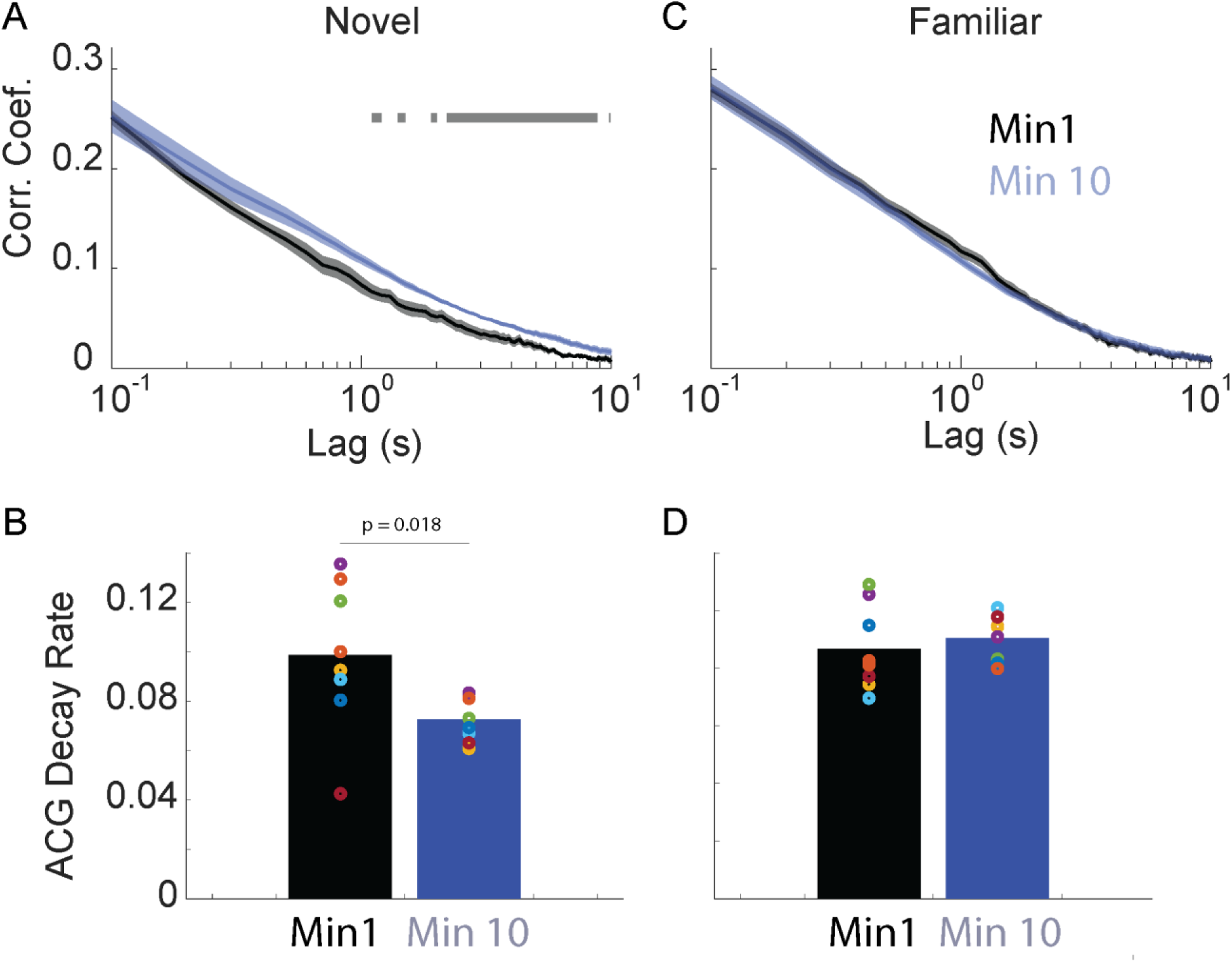
CA1 activity decorrelates faster in the first minute after transfer to a novel, but not familiar, environment. A) Population vector correction plotted as a function of lag (note log scale) during Minute 1 (black) or Minute 10 (blue) after transfer to a novel environment. Bar = p<0.01. B) The decay rate of the autocorrelation was significantly steeper in the first minute of exposure (*t(7) = 3.07, p =0.018*). C) Same as Panel A with data recorded in a familiar environment. D) No difference in ACG decay rates during the minute 1 vs minute 10 of exposure to a familiar environment (*t(7) = 0.50, p =0.63*).

**Figure S7.**
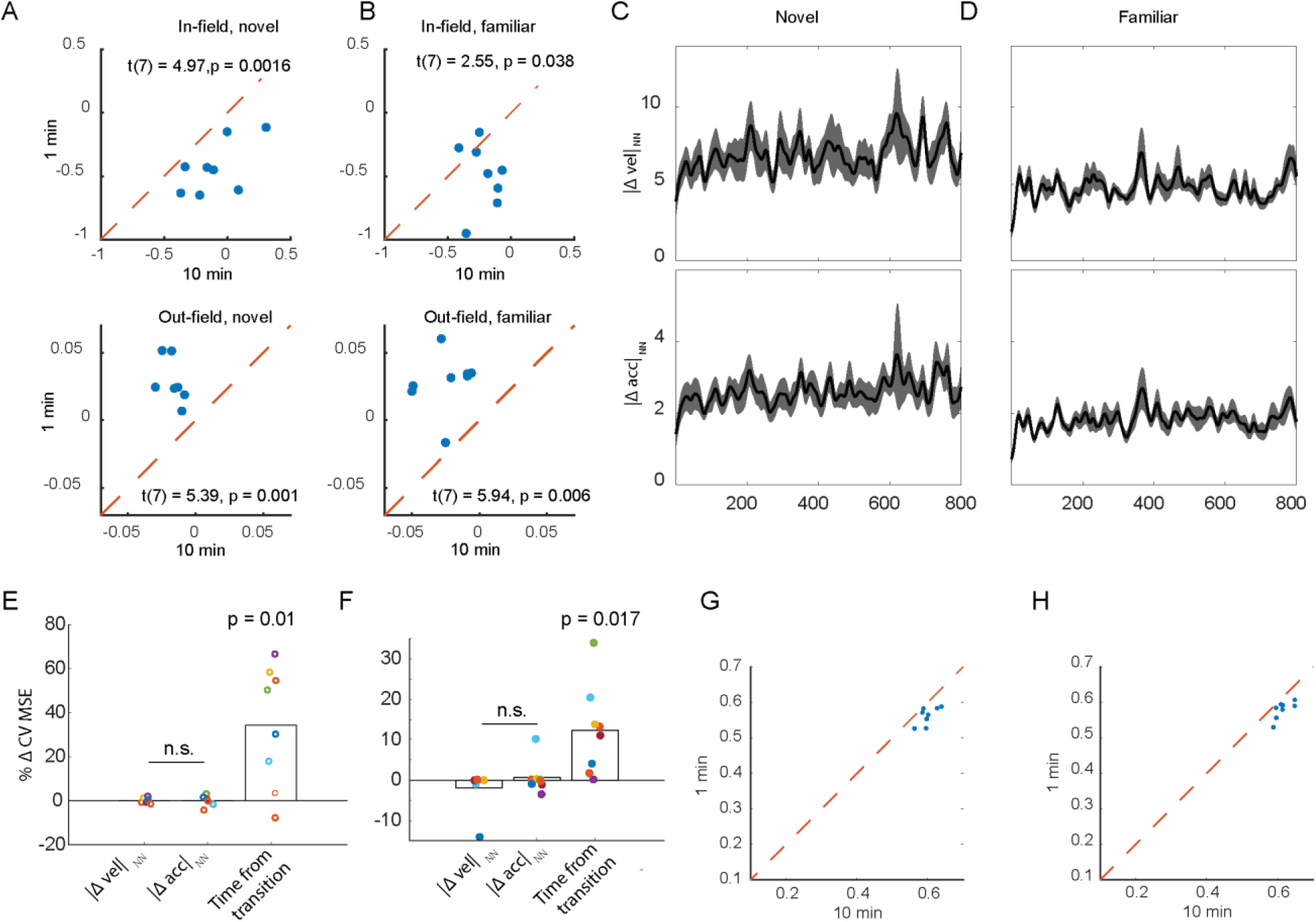
Variations in velocity and acceleration do not explain time-dependent changes in nearest-neighbor (NN) representational similarity. A) Deviations in z-scored firing rates from the mean place field activity in a novel environment. *Top,* firing rates within a place field increased over time. *Bottom*, out-of-field firing decreased over time. B) Same as Panel A with data recorded in familiar environments. C) At each moment after transitioning to a novel environment, we identified another 100-ms time bin with the most similar neural representational and calculated the absolute difference in velocity (|Δvel|_NN_) and acceleration (|Δacc|_NN_) recorded at these times. As compared to Figure 7C, neither |Δvel|_NN_ nor |Δacc|_NN_ co-varies with time as did the measure of representational uniqueness. D) Same as Panel C recorded after a transition to a familiar environment. E) Only removing time from transition decreased ability to predict NN representational similarity*, t(7) = 3.52, p = 0.01*. F) Same as panel E, recorded in a familiar environment, *t(7) = 3.12, p = 0.017*. G) In a novel environment, the patterns recorded in the first minute were less correlated to others captured in the same recording session than those observed 10 minutes into the recording (*t(7) = 5.23, p = 0.001*). H) Same as Panel G recorded in a familiar environment (t*(7) = 5.60, p = 0.0008*).

## References

1 Clewett, D., Gasser, C. & Davachi, L. Pupil-linked arousal signals track the temporal organization of events in memory. Nat Commun 11, 4007, doi:10.1038/s41467-020-17851-9 (2020).

2 Hagena, H., Hansen, N. & Manahan-Vaughan, D. beta-Adrenergic Control of Hippocampal Function: Subserving the Choreography of Synaptic Information Storage and Memory. Cereb Cortex 26, 1349–1364, doi:10.1093/cercor/bhv330 (2016).

3 Wang, S. H., Redondo, R. L. & Morris, R. G. Relevance of synaptic tagging and capture to the persistence of long-term potentiation and everyday spatial memory. Proc Natl Acad Sci U S A 107, 19537–19542, doi:10.1073/pnas.1008638107 (2010).

4 Lisman, J. E. & Grace, A. A. The hippocampal-VTA loop: controlling the entry of information into long-term memory. Neuron 46, 703–713, doi:10.1016/j.neuron.2005.05.002 (2005).

5 Sajikumar, S. & Frey, J. U. Late-associativity, synaptic tagging, and the role of dopamine during LTP and LTD. Neurobiol Learn Mem 82, 12–25, doi:10.1016/j.nlm.2004.03.003 (2004).

6 Li, S. M., Cullen, W. K., Anwyl, R. & Rowan, M. J. Dopamine-dependent facilitation of LTP induction in hippocampal CA1 by exposure to spatial novelty. Nat Neurosci 6, 526–531, doi:10.1038/nn1049 (2003).

7 Otmakhova, N. A. & Lisman, J. E. D1/D5 dopamine receptors inhibit depotentiation at CA1 synapses via cAMP-dependent mechanism. J Neurosci 18, 1270–1279, doi:10.1523/JNEUROSCI.18-04-01270.1998 (1998).

8 Otmakhova, N. A. & Lisman, J. E. D1/D5 dopamine receptor activation increases the magnitude of early long-term potentiation at CA1 hippocampal synapses. J Neurosci 16, 7478–7486, doi:10.1523/JNEUROSCI.16-23-07478.1996 (1996).

9 Lemon, N., Aydin-Abidin, S., Funke, K. & Manahan-Vaughan, D. Locus Coeruleus Activation Facilitates Memory Encoding and Induces Hippocampal LTD that Depends on β-Adrenergic Receptor Activation. Cereb Cortex 19, 2827–2837, doi:10.1093/cercor/bhp065 (2009).

10 Tsetsenis, T., Badyna, J. K., Li, R. & Dani, J. A. Activation of a Locus Coeruleus to Dorsal Hippocampus Noradrenergic Circuit Facilitates Associative Learning. Front Cell Neurosci 16, 887679, doi:10.3389/fncel.2022.887679 (2022).

11 Lethbridge, R. L., Walling, S. G. & Harley, C. W. Modulation of the perforant path-evoked potential in dentate gyrus as a function of intrahippocampal β-adrenoceptor agonist concentration in urethane-anesthetized rat. Brain Behav 4, 95–103, doi:10.1002/brb3.199 (2014).

12 Bouret, S. & Sara, S. J. Network reset: a simplified overarching theory of locus coeruleus noradrenaline function. Trends Neurosci 28, 574–582, doi:10.1016/j.tins.2005.09.002 (2005).

13 Tanila, H. Noradrenergic regulation of hippocampal place cells. Hippocampus 11, 793–808, doi:10.1002/hipo.1095 (2001).

14 Dahl, D. & Winson, J. Action of norepinephrine in the dentate gyrus. I. Stimulation of locus coeruleus. Exp Brain Res 59, 491–496, doi:10.1007/BF00261339 (1985).

15 Assaf, S. Y., Mason, S. T. & Miller, J. J. Noradrenergic modulation transmission between the entorhinal cortex and the dentate gyrus of the rat [proceedings]. J Physiol 292, 52P (1979).

16 Washburn, M. & Moises, H. C. Electrophysiological correlates of presynaptic alpha 2-receptor-mediated inhibition of norepinephrine release at locus coeruleus synapses in dentate gyrus. J Neurosci 9, 2131–2140, doi:10.1523/JNEUROSCI.09-06-02131.1989 (1989).

17 Broncel, A., Bocian, R., Klos-Wojtczak, P. & Konopacki, J. Effects of locus coeruleus activation and inactivation on hippocampal formation theta rhythm in anesthetized rats. Brain Res Bull 162, 180–190, doi:10.1016/j.brainresbull.2020.05.017 (2020).

18 Lipski, W. J. & Grace, A. A. Activation and inhibition of neurons in the hippocampal ventral subiculum by norepinephrine and locus coeruleus stimulation. Neuropsychopharmacology 38, 285–292, doi:10.1038/npp.2012.157 (2013).

19 Brown, R. A., Walling, S. G., Milway, J. S. & Harley, C. W. Locus ceruleus activation suppresses feedforward interneurons and reduces beta-gamma electroencephalogram frequencies while it enhances theta frequencies in rat dentate gyrus. J Neurosci 25, 1985–1991, doi:10.1523/JNEUROSCI.4307-04.2005 (2005).

20 Harley, C. W. & Milway, J. S. Glutamate ejection in the locus coeruleus enhances the perforant path-evoked population spike in the dentate gyrus. Exp Brain Res 63, 143–150, doi:10.1007/BF00235656 (1986).

21 Harley, C. W. & Sara, S. J. Locus coeruleus bursts induced by glutamate trigger delayed perforant path spike amplitude potentiation in the dentate gyrus. Exp Brain Res 89, 581–587, doi:10.1007/BF00229883 (1992).

22 Berridge, C. W. & Foote, S. L. Effects of Locus-Coeruleus Activation on Electroencephalographic Activity in Neocortex and Hippocampus. J Neurosci 11, 3135–3145 (1991).

23 Bacon, T. J., Pickering, A. E. & Mellor, J. R. Noradrenaline Release from Locus Coeruleus Terminals in the Hippocampus Enhances Excitation-Spike Coupling in CA1 Pyramidal Neurons Via beta-Adrenoceptors. Cereb Cortex 30, 6135–6151, doi:10.1093/cercor/bhaa159 (2020).

24 Segal, M. & Bloom, F. E. The action of norepinephrine in the rat hippocampus. IV. The effects of locus coeruleus stimulation on evoked hippocampal unit activity. Brain Res 107, 513–525, doi:10.1016/0006-8993(76)90141-4 (1976).

25 Segal, M. & Bloom, F. E. The action of norepinephrine in the rat hippocampus. III. Hippocampal cellular responses to locus coeruleus stimulation in the awake rat. Brain Res 107, 499–511, doi:10.1016/0006-8993(76)90140-2 (1976).

26 Aston-Jones, G. & Cohen, J. D. Adaptive gain and the role of the locus coeruleus-norepinephrine system in optimal performance. J Comp Neurol 493, 99–110, doi:10.1002/cne.20723 (2005).

27 Pfeffer, T. et al. Catecholamines alter the intrinsic variability of cortical population activity and perception. Plos Biol 16, doi:ARTN e2003453 10.1371/journal.pbio.2003453 (2018).

28 Cremer, A., Kalbe, F., Muller, J. C., Wiedemann, K. & Schwabe, L. Disentangling the roles of dopamine and noradrenaline in the exploration-exploitation tradeoff during human decision-making. Neuropsychopharmacology 48, 1078–1086, doi:10.1038/s41386-022-01517-9 (2023).

29 Tervo, D. G. R. et al. Behavioral Variability through Stochastic Choice and Its Gating by Anterior Cingulate Cortex. Cell 159, 21–32, doi:10.1016/j.cell.2014.08.037 (2014).

30 Usher, M., Cohen, J. D., Servan-Schreiber, D., Rajkowski, J. & Aston-Jones, G. The role of locus coeruleus in the regulation of cognitive performance. Science 283, 549–554, doi:DOI 10.1126/science.283.5401.549 (1999).

31 Brown, E. et al. Simple neural networks that optimize decisions. Int J Bifurcat Chaos 15, 803–826, doi:Doi 10.1142/S0218127405012478 (2005).

32 Munn, B. R., Muller, E. J., Wainstein, G. & Shine, J. M. The ascending arousal system shapes neural dynamics to mediate awareness of cognitive states. Nat Commun 12, 6016, doi:10.1038/s41467-021-26268-x (2021).

33 Kubie, J. L. & Muller, R. U. Multiple representations in the hippocampus. Hippocampus 1, 240–242, doi:10.1002/hipo.450010305 (1991).

34 Dupret, D., O’Neill, J., Pleydell-Bouverie, B. & Csicsvari, J. The reorganization and reactivation of hippocampal maps predict spatial memory performance. Nat Neurosci 13, 995–1002, doi:10.1038/nn.2599 (2010).

35 Hollup, S. A., Molden, S., Donnett, J. G., Moser, M. B. & Moser, E. I. Accumulation of hippocampal place fields at the goal location in an annular watermaze task. J Neurosci 21, 1635–1644, doi:10.1523/JNEUROSCI.21-05-01635.2001 (2001).

36 Markus, E. J. et al. Interactions between Location and Task Affect the Spatial and Directional Firing of Hippocampal-Neurons. J Neurosci 15, 7079–7094 (1995).

37 Moita, M. A., Rosis, S., Zhou, Y., LeDoux, J. E. & Blair, H. T. Putting fear in its place: remapping of hippocampal place cells during fear conditioning. J Neurosci 24, 7015–7023, doi:10.1523/JNEUROSCI.5492-03.2004 (2004).

38 Rosenzweig, E. S., Redish, A. D., McNaughton, B. L. & Barnes, C. A. Hippocampal map realignment and spatial learning. Nat Neurosci 6, 609–615, doi:10.1038/nn1053 (2003).

39 Muller, R. U. & Kubie, J. L. The effects of changes in the environment on the spatial firing of hippocampal complex-spike cells. J Neurosci 7, 1951–1968, doi:10.1523/JNEUROSCI.07-07-01951.1987 (1987).

40 Shapiro, M. L., Tanila, H. & Eichenbaum, H. Cues that hippocampal place cells encode: Dynamic and hierarchical representation of local and distal stimuli. Hippocampus 7, 624–642, doi:Doi 10.1002/(Sici)1098-1063(1997)7:6<624::Aid-Hipo5>3.0.Co;2-E (1997).

41 Leutgeb, S. et al. Independent codes for spatial and episodic memory in hippocampal neuronal ensembles. Science 309, 619–623, doi:10.1126/science.1114037 (2005).

42 Grella, S. L. et al. Locus Coeruleus Phasic, But Not Tonic, Activation Initiates Global Remapping in a Familiar Environment. J Neurosci 39, 445–455, doi:10.1523/JNEUROSCI.1956-18.2018 (2019).

43 Silva, D., Feng, T. & Foster, D. J. Trajectory events across hippocampal place cells require previous experience. Nat Neurosci 18, 1772–1779, doi:10.1038/nn.4151 (2015).

44 Dragoi, G. & Tonegawa, S. Development of schemas revealed by prior experience and NMDA receptor knock-out. Elife 2, e01326, doi:10.7554/eLife.01326 (2013).

45 Girardeau, G., Benchenane, K., Wiener, S. I., Buzsaki, G. & Zugaro, M. B. Selective suppression of hippocampal ripples impairs spatial memory. Nat Neurosci 12, 1222–1223, doi:10.1038/nn.2384 (2009).

46 Gridchyn, I., Schoenenberger, P., O’Neill, J. & Csicsvari, J. Assembly-Specific Disruption of Hippocampal Replay Leads to Selective Memory Deficit. Neuron 106, 291–300 e296, doi:10.1016/j.neuron.2020.01.021 (2020).

47 McNamara, C. G., Tejero-Cantero, A., Trouche, S., Campo-Urriza, N. & Dupret, D. Dopaminergic neurons promote hippocampal reactivation and spatial memory persistence. Nat Neurosci 17, 1658–1660, doi:10.1038/nn.3843 (2014).

48 Singer, A. C. & Frank, L. M. Rewarded outcomes enhance reactivation of experience in the hippocampus. Neuron 64, 910–921, doi:10.1016/j.neuron.2009.11.016 (2009).

49 Nguyen, P. V. & Gelinas, J. N. Noradrenergic gating of long-lasting synaptic potentiation in the hippocampus: from neurobiology to translational biomedicine. J Neurogenet 32, 171–182, doi:10.1080/01677063.2018.1497630 (2018).

50 Abercrombie, E. D., Keller, R. W. & Zigmond, M. J. Characterization of Hippocampal Norepinephrine Release as Measured by Microdialysis Perfusion -Pharmacological and Behavioral-Studies. Neuroscience 27, 897–904, doi:Doi 10.1016/0306-4522(88)90192-3 (1988).

51 Ihalainen, J. A., Riekkinen, P., Jr. & Feenstra, M. G. Comparison of dopamine and noradrenaline release in mouse prefrontal cortex, striatum and hippocampus using microdialysis. Neurosci Lett 277, 71–74, doi:10.1016/s0304-3940(99)00840-x (1999).

52 Moreno-Castilla, P., Perez-Ortega, R., Violante-Soria, V., Balderas, I. & Bermudez-Rattoni, F. Hippocampal release of dopamine and norepinephrine encodes novel contextual information. Hippocampus 27, 547–557, doi:10.1002/hipo.22711 (2017).

53 Feng, J. et al. Monitoring norepinephrine release in vivo using next-generation GRAB(NE) sensors. Neuron, doi:10.1016/j.neuron.2024.03.001 (2024).

54 Wilson, L. R. et al. Partial or Complete Loss of Norepinephrine Differentially Alters Contextual Fear and Catecholamine Release Dynamics in Hippocampal CA1. Biol Psychiatry Glob Open Sci 4, 51–60, doi:10.1016/j.bpsgos.2023.10.001 (2024).

55 Xiang, L. et al. Behavioral correlates of activity of optogenetically identified locus coeruleus noradrenergic neurons in rats performing T-maze tasks. Sci Rep 9, 1361, doi:10.1038/s41598-018-37227-w (2019).

56 Basu, A. et al. Frontal Norepinephrine Represents a Threat Prediction Error Under Uncertainty. Biol Psychiatry, doi:10.1016/j.biopsych.2024.01.025 (2024).

57 Jordan, R. The locus coeruleus as a global model failure system. Trends in Neurosciences 47, 92–105, doi:10.1016/j.tins.2023.11.006 (2024).

58 Jordan, R. & Keller, G. B. The locus coeruleus broadcasts prediction errors across the cortex to promote sensorimotor plasticity. Elife 12, doi:10.7554/eLife.85111 (2023).

59 Foote, S. L., Astonjones, G. & Bloom, F. E. Impulse Activity of Locus Coeruleus Neurons in Awake Rats and Monkeys Is a Function of Sensory Stimulation and Arousal. P Natl Acad Sci-Biol 77, 3033–3037, doi:DOI 10.1073/pnas.77.5.3033 (1980).

60 Herve-Minvielle, A. & Sara, S. J. Rapid habituation of auditory responses of locus coeruleus cells in anaesthetized and awake rats. Neuroreport 6, 1363–1368, doi:10.1097/00001756-199507100-00001 (1995).

61 Takeuchi, T. et al. Locus coeruleus and dopaminergic consolidation of everyday memory. Nature 537, 357–362, doi:10.1038/nature19325 (2016).

62 Vankov, A., Herve-Minvielle, A. & Sara, S. J. Response to novelty and its rapid habituation in locus coeruleus neurons of the freely exploring rat. Eur J Neurosci 7, 1180–1187, doi:10.1111/j.1460-9568.1995.tb01108.x (1995).

63 Sara, S. J., Vankov, A. & Herve, A. Locus coeruleus-evoked responses in behaving rats: a clue to the role of noradrenaline in memory. Brain Res Bull 35, 457–465, doi:10.1016/0361-9230(94)90159-7 (1994).

64 Bouret, S. & Sara, S. J. Reward expectation, orientation of attention and locus coeruleus-medial frontal cortex interplay during learning. Eur J Neurosci 20, 791–802, doi:10.1111/j.1460-9568.2004.03526.x (2004).

65 Breton-Provencher, V., Drummond, G. T., Feng, J., Li, Y. & Sur, M. Spatiotemporal dynamics of noradrenaline during learned behaviour. Nature 606, 732–738, doi:10.1038/s41586-022-04782-2 (2022).

66 Su, Z. & Cohen, J. Two types of locus coeruleus norepinephrine neurons drive reinforcement learning. bioRxiv, 10.1101/2022.12.08.519670 (2022).

67 Pittaluga, A. & Raiteri, M. Release-enhancing glycine-dependent presynaptic NMDA receptors exist on noradrenergic terminals of hippocampus. Eur J Pharmacol 191, 231–234, doi:10.1016/0014-2999(90)94153-o (1990).

68 Alme, C. B. et al. Place cells in the hippocampus: eleven maps for eleven rooms. Proc Natl Acad Sci U S A 111, 18428–18435, doi:10.1073/pnas.1421056111 (2014).

69 Kaufman, A. M., Geiller, T. & Losonczy, A. A Role for the Locus Coeruleus in Hippocampal CA1 Place Cell Reorganization during Spatial Reward Learning. Neuron 105, 1018–1026 e1014, doi:10.1016/j.neuron.2019.12.029 (2020).

70 DuBrow, S. & Davachi, L. Temporal binding within and across events. Neurobiol Learn Mem 134 Pt A, 107–114, doi:10.1016/j.nlm.2016.07.011 (2016).

71 Ennaceur, A. & Delacour, J. A new one-trial test for neurobiological studies of memory in rats. 1: Behavioral data. Behav Brain Res 31, 47–59, doi:10.1016/0166-4328(88)90157-x (1988).

72 Rait, L. I., Murty, V. P. & DuBrow, S. Contextual familiarity rescues the cost of switching. Psychon Bull Rev 31, 1103–1113, doi:10.3758/s13423-023-02392-1 (2024).

73 Sara, S. J. & Segal, M. Plasticity of sensory responses of locus coeruleus neurons in the behaving rat: implications for cognition. Prog Brain Res 88, 571–585, doi:10.1016/s0079-6123(08)63835-2 (1991).

74 Huszar, R., Zhang, Y., Blockus, H. & Buzsaki, G. Preconfigured dynamics in the hippocampus are guided by embryonic birthdate and rate of neurogenesis. Nat Neurosci 25, 1201–1212, doi:10.1038/s41593-022-01138-x (2022).

75 Hill, A. J. First occurrence of hippocampal spatial firing in a new environment. Exp Neurol 62, 282–297, doi:10.1016/0014-4886(78)90058-4 (1978).

76 Frank, L. M., Stanley, G. B. & Brown, E. N. Hippocampal plasticity across multiple days of exposure to novel environments. J Neurosci 24, 7681–7689, doi:10.1523/JNEUROSCI.1958-04.2004 (2004).

77 Priestley, J. B., Bowler, J. C., Rolotti, S. V., Fusi, S. & Losonczy, A. Signatures of rapid plasticity in hippocampal CA1 representations during novel experiences. Neuron 110, 1978–1992 e1976, doi:10.1016/j.neuron.2022.03.026 (2022).

78 Mehta, M. R., Barnes, C. A. & McNaughton, B. L. Experience-dependent, asymmetric expansion of hippocampal place fields. Proc Natl Acad Sci U S A 94, 8918–8921, doi:10.1073/pnas.94.16.8918 (1997).

79 Jackson, J. & Redish, A. D. Network dynamics of hippocampal cell-assemblies resemble multiple spatial maps within single tasks. Hippocampus 17, 1209–1229, doi:10.1002/hipo.20359 (2007).

80 Wood, E. R., Dudchenko, P. A., Robitsek, R. J. & Eichenbaum, H. Hippocampal neurons encode information about different types of memory episodes occurring in the same location. Neuron 27, 623–633, doi:Doi 10.1016/S0896-6273(00)00071-4 (2000).

81 Yavich, L., Jakala, P. & Tanila, H. Noradrenaline overflow in mouse dentate gyrus following locus coeruleus and natural stimulation: real-time monitoring by in vivo voltammetry. J Neurochem 95, 641–650, doi:10.1111/j.1471-4159.2005.03390.x (2005).

82 Mitchell, K., Oke, A. F. & Adams, R. N. In vivo dynamics of norepinephrine release-reuptake in multiple terminal field regions of rat brain. J Neurochem 63, 917–926, doi:10.1046/j.1471-4159.1994.63030917.x (1994).

83 Park, J., Takmakov, P. & Wightman, R. M. In vivo comparison of norepinephrine and dopamine release in rat brain by simultaneous measurements with fast-scan cyclic voltammetry. J Neurochem 119, 932–944, doi:10.1111/j.1471-4159.2011.07494.x (2011).

84 Aston-Jones, G., Rajkowski, J. & Cohen, J. Role of locus coeruleus in attention and behavioral flexibility. Biol Psychiatry 46, 1309–1320, doi:10.1016/s0006-3223(99)00140-7 (1999).

85 Noei, S., Zouridis, I. S., Logothetis, N. K., Panzeri, S. & Totah, N. K. Distinct ensembles in the noradrenergic locus coeruleus are associated with diverse cortical states. Proc Natl Acad Sci U S A 119, e2116507119, doi:10.1073/pnas.2116507119 (2022).

86 Berridge, C. W. & Abercrombie, E. D. Relationship between locus coeruleus discharge rates and rates of norepinephrine release within neocortex as assessed by microdialysis. Neuroscience 93, 1263–1270, doi:Doi 10.1016/S0306-4522(99)00276-6 (1999).

87 Aston-Jones, G. & Bloom, F. E. Norepinephrine-containing locus coeruleus neurons in behaving rats exhibit pronounced responses to non-noxious environmental stimuli. J Neurosci 1, 887–900, doi:10.1523/JNEUROSCI.01-08-00887.1981 (1981).

88 Pittaluga, A., Bonfanti, A. & Raiteri, M. Somatostatin potentiates NMDA receptor function via activation of InsP(3) receptors and PKC leading to removal of the Mg(2+) block without depolarization. Br J Pharmacol 130, 557–566, doi:10.1038/sj.bjp.0703346 (2000).

89 Pittaluga, A., Feligioni, M., Longordo, F., Arvigo, M. & Raiteri, M. Somatostatin-induced activation and up-regulation of N-methyl-D-aspartate receptor function: mediation through calmodulin-dependent protein kinase II, phospholipase C, protein kinase C, and tyrosine kinase in hippocampal noradrenergic nerve endings. J Pharmacol Exp Ther 313, 242–249, doi:10.1124/jpet.104.079590 (2005).

90 Risso, F. et al. Nicotine exerts a permissive role on NMDA receptor function in hippocampal noradrenergic terminals. Neuropharmacology 47, 65–71, doi:10.1016/j.neuropharm.2004.02.018 (2004).

91 Henneberger, C. et al. LTP Induction Boosts Glutamate Spillover by Driving Withdrawal of Perisynaptic Astroglia. Neuron 108, 919–936 e911, doi:10.1016/j.neuron.2020.08.030 (2020).

92 Armbruster, M., Hanson, E. & Dulla, C. G. Glutamate Clearance Is Locally Modulated by Presynaptic Neuronal Activity in the Cerebral Cortex. J Neurosci 36, 10404–10415, doi:10.1523/Jneurosci.2066-16.2016 (2016).

93 Poppenk, J., Evensmoen, H. R., Moscovitch, M. & Nadel, L. Long-axis specialization of the human hippocampus. Trends Cogn Sci 17, 230–240, doi:10.1016/j.tics.2013.03.005 (2013).

94 Bright, I. M. et al. A temporal record of the past with a spectrum of time constants in the monkey entorhinal cortex. Proc Natl Acad Sci U S A 117, 20274–20283, doi:10.1073/pnas.1917197117 (2020).

95 Momennejad, I. & Howard, M. W. Predicting the future with multi-scale successor representations. bioRxiv 449470 (2018).

96 Tiganj, Z., Gershman, S. J., Sederberg, P. B. & Howard, M. W. Estimating Scale-Invariant Future in Continuous Time. Neural Comput 31, 681–709, doi:10.1162/neco_a_01171 (2019).

97 Wang, Y. C., Adcock, R. A. & Egner, T. Toward an integrative account of internal and external determinants of event segmentation. Psychon Bull Rev 31, 484–506, doi:10.3758/s13423-023-02375-2 (2024).

98 Baldassano, C. et al. Discovering Event Structure in Continuous Narrative Perception and Memory. Neuron 95, 709–721 e705, doi:10.1016/j.neuron.2017.06.041 (2017).

99 Heusser, A. C., Ezzyat, Y., Shiff, I. & Davachi, L. Perceptual Boundaries Cause Mnemonic Trade-Offs Between Local Boundary Processing and Across-Trial Associative Binding. J Exp Psychol Learn 44, 1075–1090, doi:10.1037/xlm0000503 (2018).

100 Boltz, M. Temporal accent structure and the remembering of filmed narratives. J Exp Psychol Hum Percept Perform 18, 90–105, doi:10.1037//0096-1523.18.1.90 (1992).

101 Murdock, B. B., Jr. Modality effects in short-term memory: storage or retrieval? J Exp Psychol 77, 79–86, doi:10.1037/h0025786 (1968).

102 Kesner, R. P., Chiba, A. A. & Jacksonsmith, P. Rats Do Show Primacy and Recency Effects in Memory for Lists of Spatial Locations - a Reply to Gaffan. Anim Learn Behav 22, 214–218, doi:Doi 10.3758/Bf03199922 (1994).

103 Bolhuis, J. J. & van Kampen, H. S. Serial position curves in spatial memory of rats: primacy and recency effects. Q J Exp Psychol B 40, 135–149 (1988).

104 Ezzyat, Y. & Davachi, L. Similarity breeds proximity: pattern similarity within and across contexts is related to later mnemonic judgments of temporal proximity. Neuron 81, 1179–1189, doi:10.1016/j.neuron.2014.01.042 (2014).

105 Pu, Y., Kong, X. Z., Ranganath, C. & Melloni, L. Event boundaries shape temporal organization of memory by resetting temporal context. Nat Commun 13, 622, doi:10.1038/s41467-022-28216-9 (2022).

106 Sinclair, A. H., Manalili, G. M., Brunec, I. K., Adcock, R. A. & Barense, M. D. Prediction errors disrupt hippocampal representations and update episodic memories. Proc Natl Acad Sci U S A 118, doi:10.1073/pnas.2117625118 (2021).

107 Kim, G., Norman, K. A. & Turk-Browne, N. B. Neural Differentiation of Incorrectly Predicted Memories. J Neurosci 37, 2022–2031, doi:10.1523/JNEUROSCI.3272-16.2017 (2017).

108 Carter, M. E. et al. Tuning arousal with optogenetic modulation of locus coeruleus neurons. Nat Neurosci 13, 1526–1533, doi:10.1038/nn.2682 (2010).

109 Murphy, P. R., O’Connell, R. G., O’Sullivan, M., Robertson, I. H. & Balsters, J. H. Pupil diameter covaries with BOLD activity in human locus coeruleus. Hum Brain Mapp 35, 4140–4154, doi:10.1002/hbm.22466 (2014).

110 Megemont, M., McBurney-Lin, J. & Yang, H. D. Pupil diameter is not an accurate real-time readout of locus coeruleus activity. Elife 11 (2022).

111 Wilmot, J. H. et al. Phasic locus coeruleus activity enhances trace fear conditioning by increasing dopamine release in the hippocampus. Elife 12, doi:10.7554/eLife.91465 (2024).

112 Tse, D. et al. Cell-type-specific optogenetic stimulation of the locus coeruleus induces slow-onset potentiation and enhances everyday memory in rats. Proc Natl Acad Sci U S A 120, e2307275120, doi:10.1073/pnas.2307275120 (2023).

113 Chowdhury, A. et al. A locus coeruleus-dorsal CA1 dopaminergic circuit modulates memory linking. Neuron 110, 3374–3388 e3378, doi:10.1016/j.neuron.2022.08.001 (2022).

114 Wagatsuma, A. et al. Locus coeruleus input to hippocampal CA3 drives single-trial learning of a novel context. Proc Natl Acad Sci U S A 115, E310–E316, doi:10.1073/pnas.1714082115 (2018).

115 Kempadoo, K. A., Mosharov, E. V., Choi, S. J., Sulzer, D. & Kandel, E. R. Dopamine release from the locus coeruleus to the dorsal hippocampus promotes spatial learning and memory. Proc Natl Acad Sci U S A 113, 14835–14840, doi:10.1073/pnas.1616515114 (2016).

116 Sara, S. J. Locus Coeruleus in time with the making of memories. Curr Opin Neurobiol 35, 87–94, doi:10.1016/j.conb.2015.07.004 (2015).

117 Hämmerer, D. et al. Locus coeruleus integrity in old age is selectively related to memories linked with salient negative events. P Natl Acad Sci USA 115, 2228–2233, doi:10.1073/pnas.1712268115 (2018).

118 Seo, D. O. et al. A locus coeruleus to dentate gyrus noradrenergic circuit modulates aversive contextual processing. Neuron 109, 2116–2130 e2116s, doi:10.1016/j.neuron.2021.05.006 (2021).

119 Amaral, D. G. & Foss, J. A. Locus Coeruleus Lesions and Learning. Science 188, 377–378, doi:DOI 10.1126/science.1118734 (1975).

120 Frey, U. & Morris, R. G. M. Synaptic tagging and long-term potentiation. Nature 385, 533–536, doi:DOI 10.1038/385533a0 (1997).

121 O’Carroll, C. M., Martin, S. J., Sandin, J., Frenguelli, B. & Morris, R. G. Dopaminergic modulation of the persistence of one-trial hippocampus-dependent memory. Learn Mem 13, 760–769, doi:10.1101/lm.321006 (2006).

122 He, K. W. et al. Distinct Eligibility Traces for LTP and LTD in Cortical Synapses. Neuron 88, 528–538, doi:10.1016/j.neuron.2015.09.037 (2015).

123 Harley, C., Milway, J. S. & Lacaille, J. C. Locus coeruleus potentiation of dentate gyrus responses: evidence for two systems. Brain Res Bull 22, 643–650, doi:10.1016/0361-9230(89)90084-1 (1989).

124 Frey, S., Bergado-Rosado, J., Seidenbecher, T., Pape, H. C. & Frey, J. U. Reinforcement of early long-term potentiation (early-LTP) in dentate gyrus by stimulation of the basolateral amygdala: heterosynaptic induction mechanisms of late-LTP. J Neurosci 21, 3697–3703, doi:10.1523/JNEUROSCI.21-10-03697.2001 (2001).

125 Pignatelli, M. et al. Engram Cell Excitability State Determines the Efficacy of Memory Retrieval. Neuron 101, 274–284 e275, doi:10.1016/j.neuron.2018.11.029 (2019).

126 Meenakshi, P., Kumar, S. & Balaji, J. In vivo imaging of immediate early gene expression dynamics segregates neuronal ensemble of memories of dual events. Mol Brain 14, 102, doi:10.1186/s13041-021-00798-3 (2021).

127 Cai, D. J. et al. A shared neural ensemble links distinct contextual memories encoded close in time. Nature 534, 115–118, doi:10.1038/nature17955 (2016).

128 Yang, W. et al. Selection of experience for memory by hippocampal sharp wave ripples. Science 383, 1478–1483, doi:10.1126/science.adk8261 (2024).

129 Jezek, K., Henriksen, E. J., Treves, A., Moser, E. I. & Moser, M. B. Theta-paced flickering between place-cell maps in the hippocampus. Nature 478, 246–249, doi:10.1038/nature10439 (2011).

130 Chung, A. et al. Cognitive control persistently enhances hippocampal information processing. Nature 600, 484–488, doi:10.1038/s41586-021-04070-5 (2021).

131 Pettit, N. L., Yuan, X. C. & Harvey, C. D. Hippocampal place codes are gated by behavioral engagement. Nat Neurosci 25, 561–566, doi:10.1038/s41593-022-01050-4 (2022).

132 El-Gaby, M. et al. An emergent neural coactivity code for dynamic memory. Nat Neurosci 24, 694–704, doi:10.1038/s41593-021-00820-w (2021).

133 Monaco, J. D., Rao, G., Roth, E. D. & Knierim, J. J. Attentive scanning behavior drives one-trial potentiation of hippocampal place fields. Nat Neurosci 17, 725–731, doi:10.1038/nn.3687 (2014).

134 Olpe, H. R. et al. Glutamate-Induced Activation of Rat Locus-Coeruleus Increases Ca1 Pyramidal Cell Excitability. Neurosci Lett 65, 11–16, doi:Doi 10.1016/0304-3940(86)90112-6 (1986).

135 Kafkas, A. & Montaldi, D. How do memory systems detect and respond to novelty? Neurosci Lett 680, 60–68, doi:10.1016/j.neulet.2018.01.053 (2018).

136 Ben-Yakov, A. & Henson, R. N. The Hippocampal Film Editor: Sensitivity and Specificity to Event Boundaries in Continuous Experience. J Neurosci 38, 10057–10068, doi:10.1523/JNEUROSCI.0524-18.2018 (2018).

137 VanElzakker, M., Fevurly, R. D., Breindel, T. & Spencer, R. L. Environmental novelty is associated with a selective increase in Fos expression in the output elements of the hippocampal formation and the perirhinal cortex. Learn Memory 15, 899–908, doi:10.1101/lm.1196508 (2008).

138 Larkin, M. C., Lykken, C., Tye, L. D., Wickelgren, J. G. & Frank, L. M. Hippocampal output area CA1 broadcasts a generalized novelty signal during an object-place recognition task. Hippocampus 24, 773–783, doi:10.1002/hipo.22268 (2014).

139 Jenkins, T. A., Amin, E., Pearce, J. M., Brown, M. W. & Aggleton, J. P. Novel spatial arrangements of familiar visual stimuli promote activity in the rat hippocampal formation but not the parahippocampal cortices: a c-fos expression study. Neuroscience 124, 43–52, doi:10.1016/j.neuroscience.2003.11.024 (2004).

140 Allen, T. A., Salz, D. M., McKenzie, S. & Fortin, N. J. Nonspatial Sequence Coding in CA1 Neurons. J Neurosci 36, 1547–1563, doi:10.1523/JNEUROSCI.2874-15.2016 (2016).

141 Vinogradova, O. S. Hippocampus as comparator: role of the two input and two output systems of the hippocampus in selection and registration of information. Hippocampus 11, 578–598, doi:10.1002/hipo.1073 (2001).

142 Kumaran, D. & Maguire, E. A. An unexpected sequence of events: mismatch detection in the human hippocampus. Plos Biol 4, e424, doi:10.1371/journal.pbio.0040424 (2006).

143 Aston-Jones, G. et al. Afferent regulation of locus coeruleus neurons: anatomy, physiology and pharmacology. Prog Brain Res 88, 47–75, doi:10.1016/s0079-6123(08)63799-1 (1991).

144 Kishi, T. et al. Topographical organization of projections from the subiculum to the hypothalamus in the rat. Journal of Comparative Neurology 419, 205–222, doi:Doi 10.1002/(Sici)1096-9861(20000403)419:2<05::Aid-Cne5>3.0.Co;2-0 (2000).

145 Rolls, E. T. in Neural models of plasticity Ch. 13, 240–265 (Academic Press, 1989).

146 Oleskevich, S., Descarries, L. & Lacaille, J. C. Quantified distribution of the noradrenaline innervation in the hippocampus of adult rat. J Neurosci 9, 3803–3815, doi:10.1523/JNEUROSCI.09-11-03803.1989 (1989).

147 Ryan, J. & Rogers, M. Event Segmentation Deficits in ADHD. J Atten Disord 25, 355–363, doi:10.1177/1087054718799929 (2021).

148 Zalla, T., Verlut, I., Franck, N., Puzenat, D. & Sirigu, A. Perception of dynamic action in patients with schizophrenia. Psychiatry Res 128, 39–51, doi:10.1016/j.psychres.2003.12.026 (2004).

149 Zacks, J. M., Speer, N. K., Vettel, J. M. & Jacoby, L. L. Event understanding and memory in healthy aging and dementia of the Alzheimer type. Psychol Aging 21, 466–482, doi:10.1037/0882-7974.21.3.466 (2006).

150 Braak, H., Thal, D. R., Ghebremedhin, E. & Del Tredici, K. Stages of the pathologic process in Alzheimer disease: age categories from 1 to 100 years. J Neuropathol Exp Neurol 70, 960–969, doi:10.1097/NEN.0b013e318232a379 (2011).

151 Theofilas, P. et al. Locus coeruleus volume and cell population changes during Alzheimer’s disease progression: A stereological study in human postmortem brains with potential implication for early-stage biomarker discovery. Alzheimers Dement 13, 236–246, doi:10.1016/j.jalz.2016.06.2362 (2017).

152 Nisenbaum, L. K., Zigmond, M. J., Sved, A. F. & Abercrombie, E. D. Prior exposure to chronic stress results in enhanced synthesis and release of hippocampal norepinephrine in response to a novel stressor. J Neurosci 11, 1478–1484, doi:10.1523/JNEUROSCI.11-05-01478.1991 (1991).

153 Belujon, P. & Grace, A. A. Hippocampus, amygdala, and stress: interacting systems that affect susceptibility to addiction. Ann Ny Acad Sci 1216, 114–121, doi:10.1111/j.1749-6632.2010.05896.x (2011).

154 Simpson, E. H. et al. Lights, fiber, action! A primer on in vivo fiber photometry. Neuron 112, 718–739, doi:10.1016/j.neuron.2023.11.016 (2024).

155 Keevers, L. J., McNally, G. P. & Jean-Richard-dit-Bressel, P. Obtaining artifact-corrected signals in fiber photometry: Isosbestic signals, robust regression and dF/F calculations. Research Square Platform, 10.21203/rs.3.rs-3549461/v1 (2023).

156 Mathis, A. et al. DeepLabCut: markerless pose estimation of user-defined body parts with deep learning. Nat Neurosci 21, 1281–1289, doi:10.1038/s41593-018-0209-y (2018).

157 Petersen, P. C., Hernandez, M. & Buzsáki, G. (Zenodo, 2020).

158 Pachitariu, M., Steinmetz, N., Kadir, S., Carandini, M. & Harris, K. Fast and accurate spike sorting of high-channel count probes with KiloSort. Adv Neur In 29 (2016).

159 Rossant, C. et al. Spike sorting for large, dense electrode arrays. Nat Neurosci 19, 634–641, doi:10.1038/nn.4268 (2016).

